# Machine learning-based meta-analysis reveals gut microbiome alterations associated with Parkinson’s disease

**DOI:** 10.1101/2023.12.05.569565

**Authors:** Stefano Romano, Jakob Wirbel, Rebecca Ansorge, Christian Schudoma, Quinten Raymond Ducarmon, Arjan Narbad, Georg Zeller

**Author notes:** Department of Medical Microbiology and Leiden University Center for Infectious Diseases, Leiden University Medical Center, Leiden, Netherlands (Georg Zeller).

## Abstract

There is strong interest in exploring the potential of the gut microbiome for Parkinson’s disease (PD) diagnosis and treatment. However, a consensus on the microbiome features associated with PD and a multi-study assessment of their diagnostic value is lacking. Here, we present a machine learning meta-analysis of PD microbiome studies of unprecedented scale (including 4,490 samples). Within most studies, microbiome-based machine learning models could accurately classify PD patients. However, models were study-specific and did not generalise well across other studies. By training models on multiple datasets, we could improve their general applicability and disease specificity as assessed against microbiomes from other neurodegenerative diseases. Meta-analysis of shotgun metagenomes moreover delineated PD-associated microbial pathways potentially contributing to the deterioration of gut health and favouring the translocation of pathogenic molecules along the gut-brain axis. Strikingly, diverse microbial pathways for the biotransformation of solvents and pesticides were enriched in PD. These results align with the epidemiological evidence that exposure to these molecules increases PD risk and raise the question of whether gut microbial metabolism modulates their toxicity. Taken together, we offer the most comprehensive overview to date about the PD gut microbiome and provide future reference for its diagnostic and functional potential.

## Introduction

Parkinson’s disease (PD) is the second most common age-related neurodegenerative disease after Alzheimer’s disease. Recent estimates suggest a doubling of PD patients every ∼30 years, which might result in around 12 million patients worldwide by 2050^1^. Only a minority of PD cases are thought to be of purely genetic origin and environmental factors are of crucial importance in disease development^2–4^. A hallmark of PD is the accumulation of Lewy’s bodies containing misfolded α-synuclein (αSyn) proteins in the central nervous system (CNS), causing neuron toxicity and death^5^. Specifically, the loss of dopaminergic neurons and consequent decrease in dopamine levels are the molecular mechanisms underlying motor impairments observed in PD patients^5^. However, PD manifests with a plethora of both motor and non-motor symptoms, many of which involve the gastrointestinal (GI) tract ^6–8^. Among the latter, gastroparesis, gut inflammation, increased intestinal permeability, and constipation are frequently observed^8^ and some of these GI symptoms have been shown to be predictive of PD^7^. Surprisingly, the GI tract involvement can precede motor symptoms by many years. For example, constipation is among the earliest non-motor symptoms and can appear up to twenty years before diagnosis^9^. Moreover, recent evidence has linked GI inflammatory diseases, such as IBD, to PD pathophysiology^10,11^. This relationship between GI health and PD has motivated numerous investigations of the putative roles of the gut microbiome in the disease.

We recently conducted a meta-analysis of gut microbiome studies in PD (based on 16S ribosomal RNA gene amplicon sequencing) and showed that when compared to controls, the gut microbiome of PD patients has some common alterations across patient populations from diverse countries and continents^12^. Although high variability between studies was observed, as often in microbiome meta-analyses^13^, the gut microbiome in PD patients is typically depleted in short-chain fatty acid (SCFA) producing bacteria. SCFAs are the end product of the bacterial fermentation of complex carbohydrates and they play a pivotal role in maintaining epithelial barrier integrity and colonic immune homeostasis. Similar results have been confirmed by independent meta-analyses and more recent shotgun metagenomic studies^14–16^. Nevertheless, there is still limited consensus on the bacterial species and metabolic pathways associated with the disease^15–19^. Identifying microbial taxa and especially metabolic functions associated with PD across sampling populations is essential in order to develop mechanistic hypotheses through which the microbiome could possibly contribute to the disease. This will open doors for 1) designing *in-vitro*/*in-vivo* experiments to mechanistically elucidate a putative impact of gut microbes on PD; 2) developing tools and strategies to use the microbiome for disease diagnosis, prognosis, and treatment.

To date PD is diagnosed through clinical assessment of motor symptoms, which appear when between 50 and 80% of dopaminergic neurons have already been lost^5^, hampering the possibilities of testing neuroprotective therapies. Hence, there is a strong interest in identifying potential markers to foster early diagnosis. To address this, several attempts have been made to use gut microbiome features for building machine learning (ML) classification models that discriminate PD patients from controls^17,18,20–22^, reporting up to 90% classification accuracies. However, we currently do not know whether these prediction accuracies are observed across multiple datasets conducted in different countries. Specifically, model portability, indicating how well models perform when applied to an independent dataset obtained from another sampling population, has never been investigated in the context of PD. This is, however, relevant as it could reveal features (i.e. bacterial taxa/functions) that consistently discriminate PD from controls, thus informing on the potential generalisation and global applicability of such models. Finally, the combination of multiple datasets in a large-scale meta-analysis could ideally lead to more accurate and robust models for PD classification, and it has so far not been thoroughly explored.

To fill this knowledge gap, here we perform a large-scale meta-analysis of the gut microbiome in PD to assess how accurately ML models based on the currently available gut microbiome data can discriminate PD patients from controls. We use both public 16S amplicon sequencing and shotgun metagenomics data to extensively evaluate various ML approaches based on single and combined datasets. We complement this ML analysis by conducting the largest meta-analysis so far on the gut microbiome in PD to establish an updated list of prokaryotic taxa and microbial metabolic functions robustly associated with the disease.

**Table 1.** Overview of the studies used in this work. Samples refer to those used in our study after data filtration. CTRL = controls; PD = Parkinson’s disease patients. ENA = £; GSA = $; Figshare = &; DDBJ = *. 16S = 16S ribosomal RNA gene amplicon data; SMG = shotgun metagenomic data.

| Study | Accession Id | Data type | #Samples | Country |
| --- | --- | --- | --- | --- |
| Wallen 2021<sup>23</sup> (2 datasets) | PRJNA601994<sup>£</sup> | 16S | 130/184 CTRL; 196/323 PD | USA |
| Zhang 2020<sup>24</sup> | CRA001938<sup>\$</sup> | 16S | 137 CTRL; 63 PD | China |
| Tan 2021<sup>25</sup> | PRJNA494620<sup>£</sup> | 16S | 96 CTRL; 104 PD | Malaysia |
| Heintz-Buschart 2017<sup>26</sup> | PRJNA381395<sup>£</sup> | 16S | 38 CTRL; 26 PD | Germany |
| Petrov 2017<sup>27</sup> | *Personal communication* | 16S | 67 CTRL; 95 PD | Russia |
| Qian 2018<sup>28</sup> | PRJNA391524<sup>£</sup> | 16S | 45 CTRL; 45 PD | China |
| Keshavarzian 2015<sup>29</sup> | PRJNA268515<sup>£</sup> | 16S | 31 CTRL; 34 PD | USA |
| Pietrucci 2019<sup>30</sup> | PRJNA510730<sup>£</sup> | 16S | 72 CTRL; 80 PD | Italy |
| Aho 2019<sup>31</sup> | PRJEB27564<sup>£</sup> | 16S | 65 CTRL; 67 PD | Finland |
| Weis 2019<sup>32</sup> | PRJEB30615<sup>£</sup> | 16S | 25 CTRL; 39 PD | Germany |
| Hopfner 2017<sup>33</sup> | PRJEB14928<sup>£</sup> | 16S | 26 CTRL; 29 PD | Germany |
| Nishiwaki 2020<sup>14</sup> | DRA009229* | 16S | 137 CTRL; 223 PD | Japan |
| Cirstea 2020<sup>34</sup> | *Personal communication* | 16S | 103 CTRL; 197 PD | Canada |
| Lubomski 2022<sup>21</sup> | PRJNA808166<sup>£</sup> | 16S | 81 CTRL; 103 PD | Australia |
| Kenna 2021<sup>35</sup> | DOI:<br>[10.6084/m9.figshare.14345513](https://doi.org/10.6084/m9.figshare.14345513)<sup>&</sup> | 16S | 47 CTRL; 86 PD | Canada |
| Jo 2022<sup>19</sup> | PRJNA742875<sup>£</sup> | 16S | 84 CTRL; 88 PD | China |
| **Total samples** |  | **16S** | **HC 1368; PD 1798** |  |
| Bedarf 2017<sup>18</sup> | PRJEB17784<sup>£</sup> | SMG | 28 CTRL; 31 PD | Germany |
| Qian 2020<sup>17</sup> | PRJNA433459<sup>£</sup> | SMG | 40 CTRL; 40 PD | China |
| Mao 2021<sup>36</sup> | PRJNA588035<sup>£</sup> | SMG | 39 CTRL; 39 PD | China |
| Jo 2022<sup>19</sup> | PRJNA743718<sup>£</sup> | SMG | 74 CTRL; 82 PD | China |
| Wallen 2022<sup>15</sup> | PRJNA834801<sup>£</sup> | SMG | 234 CTRL; 490 PD | USA |
| Boktor 2023<sup>16</sup> (2 datasets) | ERP138197/ERP138199<sup>£</sup> | SMG | 72/67 CTRL; 42/46 PD | USA |
| **Total samples** |  | **SMG** | **554 HC; 770 PD** |  |

## Results

### Datasets overview and beta diversity analysis

We processed a total of 4,490 samples obtained from case-control studies across 11 countries and 4 continents that profiled the faecal microbiome of PD patients and controls using 16S amplicon (16S; 3,166 samples) and shotgun metagenomics sequencing (SMG; 1,324 samples; Table 1). Altogether, the number of samples we used in this study is up to four times larger than those used in previous PD meta-analyses^12,14,16,22^. This first allowed us to investigate the overall structure of the microbiome through a well-powered beta diversity analysis. Consistent with our previous report^12^, samples did not cluster according to disease status in ordination plots visualising beta diversity (Fig 1a, 1b). Even after removing the effect of the study of origin, only a weak separation was observed between PD and controls (Fig. 1c, 1d). Permutational multivariate analysis of variance (Permanova) indicated that the disease status explains ≤1% of the variance in microbiome composition across studies, yet this was statistically highly significant (Fig 1). The study of origin instead explains a considerably higher proportion of the data, 19.9% and 7.7% for the 16S and SMG data, respectively. This highlights the strong variability in microbiome composition across studies that is often observed in microbiome meta-analyses^12,13^.

### Comparison of machine learning approaches

To assess how well the microbiome profiles could distinguish between control and PD samples, we first applied ML models to each dataset individually. Since any ML workflow requires a multitude of parameter choices, we initially explored and compared different filtering strategies, normalisation approaches, and ML algorithms implemented in the R package SIAMCAT v_2.0^37^. Accuracies of ML models were evaluated using the area under the receiver operating characteristics curve (AUC). These comparisons were performed for the taxonomies of both 16S and SMG data. For both types of data, retaining taxa detected in at least 5% of the samples in 10 16S and 2 SMG datasets resulted in profiles which allowed to build ML models with the highest AUCs (Fig S1, S2). However, the accuracies of models varied greatly across ML algorithms and filtering/normalisation strategies (Fig S1, S2). For the 16S data, Random Forest and Ridge regression (including the LibLinear implementation) classifiers performed considerably better than the other algorithms tested, reaching a maximum AUC of ≥95%, observed in within-study cross-validation (CV) performed for the data of Zhang et al.^24^ and Tan et al.^25^ (Fig S3). Similarly variable results were obtained for the SMG data, for which the LASSO algorithm (LibLinear implementation) and the Ridge regression yielded the most accurate models with 90 and 88% AUCs in the study of Bedarf et al.^18^ and Qian et al^17^ (Fig S4). For the sake of clarity and comparability, all results presented in the main text were obtained using the Ridge regression classifiers for both 16S and SMG data (Fig 2). Between the two data types, ML models built on SMG data had a higher average AUC for the within-study CV and a considerably lower variation compared to the 16S-based models (Fig 2; SMG = 78.3% ± 6.50, 16S = 72.3% ± 11.7; two samples Welch t-test: t = -1.6, df = 19.6, *p-value* = 0.13). For both data types no correlation was observed between the number of samples used to train the models and classification accuracies (Pearson correlation, 16S: t = 0.27, df = 15, p-value = 0.8; SMG: t = 0.1, df = 5, p-value = 0.93).

### Cross-study portability of the ML models

To verify portability of the ML models, which refers to the classification accuracies of a study-specific model when tested on an independent holdout study, we performed a study-to-study validation (cross study validation; CSV) for both 16S and SMG data. In this approach, the models built for each dataset were tested on all other datasets of the same data type. Compared to the performance estimated through within-study CV, CSV performances were significantly lower for both SMG and 16S data (two samples Welch t-test: SMG t = -5.2, df = 8.8, p-value < 0.001, 16S t = -3.7, df = 16.9, p-value < 0.02; Fig 2, S3, S4). In general, 16S datasets showing high AUC in the within-study CV (i.e. Petrov et al.^27^, Zhang et al.^24^, and Tan et al.^25^) could also be better classified by external models (i.e. models built on other datasets; Fig S4). However, the models trained on these datasets showed a much lower performance when tested on hold-out data (Fig S4). For example, the Ridge model built on the data from Petrov et al.^27^ had an AUC of 88% in within-study CV, but its average AUC on the other datasets was as low as 56.9%. A similar pattern was observed for the SMG data, where models had a higher overall average performance in the within-study CV, and performed substantially worse in the CSV (Fig 2, S5). For example, the datasets of Bedarf et al.^18^ resulted in a Ridge model with a within-study CV AUC of 85% in contrast to an average CSV AUC of only 57.4% (Fig S4). CSV AUCs obtained for the SMG models were higher than those obtained for the 16S models (two samples Welch t-test: t = -2.1, df = 55.97, p-value = 0.04; average AUC SMG 64.2% ± 7.42, average AUC 16S 61.6% ± 7.81). As for the CV, for both data types no correlation was observed between the number of samples used for training and the CSV classification accuracies (Pearson correlation, 16S: t = -0.14, df = 270, p-value = 0.9; SMG: t = 1.22, df = 40, p-value = 0.23). These low overall CSV performances observed in both the 16S and SMG data indicate large inter-study variability in microbiome composition, consistent with the Permanova results (Fig. 1).

**Fig 1.**
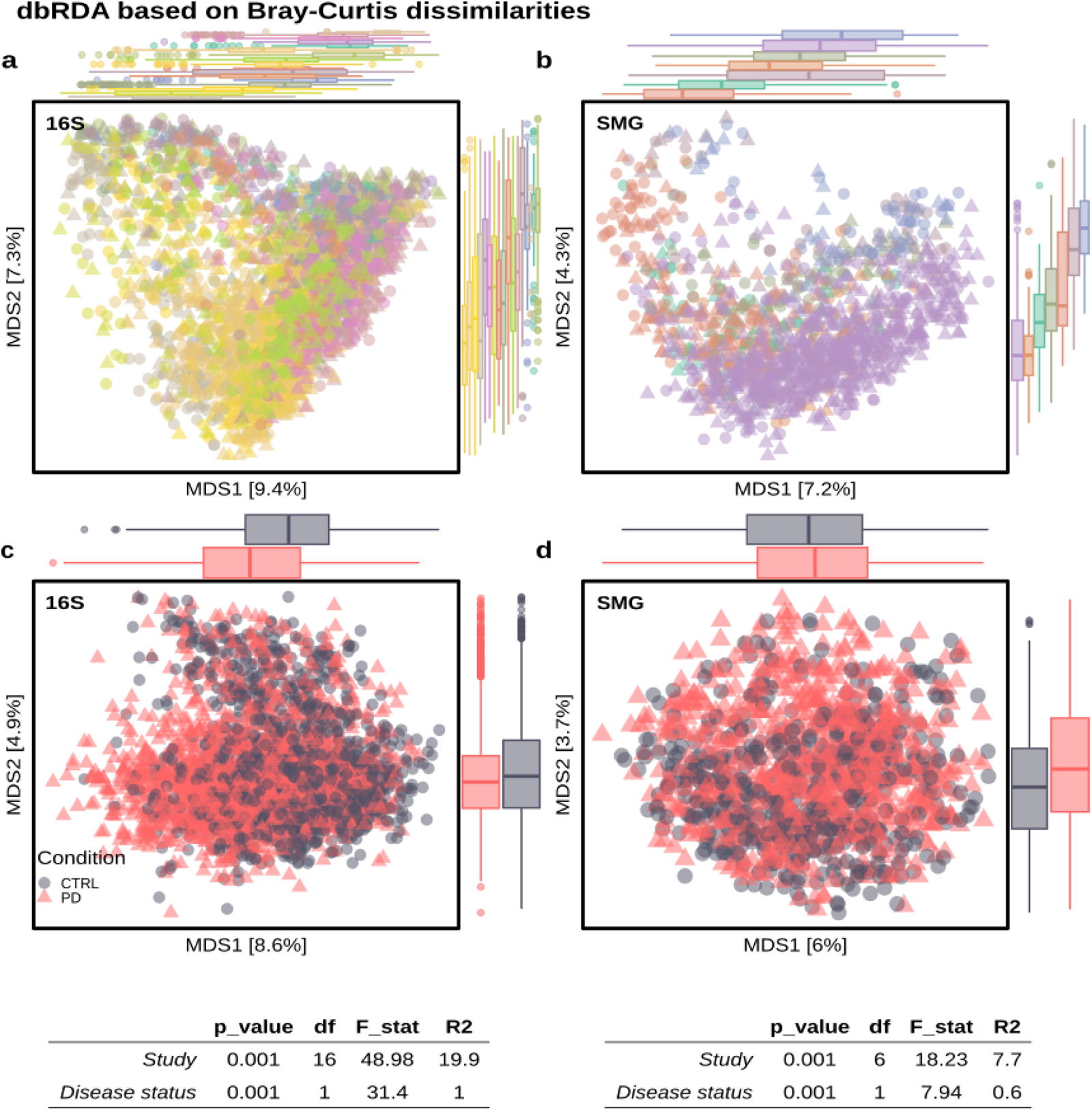
The composition of the gut microbiome is significantly different between PD and controls (CTRL). Distance-based redundancy analysis (dbRDA) was performed on Bray-Curtis dissimilarities calculated for the 16S (a, c) and SMG data (b, d). Raw distances (a, b) and distances conditioned by the study of origin to remove study-specific batch effects are shown (c, d). Side boxplots depict the sample distribution along the two components and are coloured to indicate the different datasets used (a, b) and the disease status (c, d). Edges of the boxplots indicate the first and third quartile (interquartile range, IQR), and the middle thick segment indicates the sample median. Side whiskers extend to the largest and smallest values till 1.5 x IQR. Data beyond this range are reported as dots. Results of permutational analysis of variance (Permanova) performed for each data type are shown in the bottom tables. The significance of the clustering was calculated for both the study of origin and the disease status. For the latter, the permutations were restricted within the study of origin. Legend keys for the colours in panel a and b are reported in Fig S8.

The re-analyzed 16S datasets varied greatly in sequencing depths (Fig S5). Heterogeneous sequencing depth may negatively affect the generalisation capabilities of ML models. To test if model performances were impacted by sequencing depth, we built ML models on rarefied data (2000 read counts final depth), and compared their accuracy in CSV to those of the not-rarefied models. For the majority of the ML algorithms, sequencing depth did not noticeably influence the model performances and AUC did not change significantly between the two approaches (paired Welch t-test: ENET t = -0.2, df = 271, p-value = 0.82; LASSO t = -0.5, df = 271, p-value = 0.59; LASSO-LibLinear t = -0.9, df = 271, p-value = 0.35; Random Forest t = 0.4, df = 271, p-value = 0.67; Ridge t = -1.96, df = 271, p-value = 0.05; average difference in AUCs < 0.7%; Fig S6). Only Ridge regression (as implemented in LibLinear) was sensitive to heterogeneity in sequencing depth (paired Welch t-test: Ridge-LibLinear t = -4.4, df 271, p-value <0.001; average AUC rarefied data = 62.3% vs average AUC not rarefied data 60.9%). Another factor potentially affecting CSV performances is study-associated heterogeneity in microbiome profiles – due to technical or biological differences – also referred to as batch effects (here rather study effects). As our beta-diversity analysis suggests batch effects to be considerably higher in the 16S data than the SMG data (Fig 1), we investigated if correcting for batch effects in the 16S data would increase the overall model portability. We used various available batch correction approaches and, to avoid overoptimistic evaluations^38^, we ensured that all methods were blind to the labels (PD vs controls). However, none of the batch correction approaches used here significantly increased the average AUC in the CSV evaluations (ANOVA: ENET F = 0.3, df = 4, p-value = 0.87; LASSO F = 0.2, df = 4, p-value = 0.95; LASSO-LibLinear F = 0.41, df = 4, p-value = 0.8; Random Forest F = 0.002, df = 4, p-value = 1; Ridge F = 0.2, df = 4, p-value = 0.9; Ridge-LibLinear F = 0.09, df = 4, p-value = 0.98; Fig S7). This suggests that currently available batch correction methods may be of limited practical value for improving cross-study portability of microbiome classifiers.

**Fig 2.**
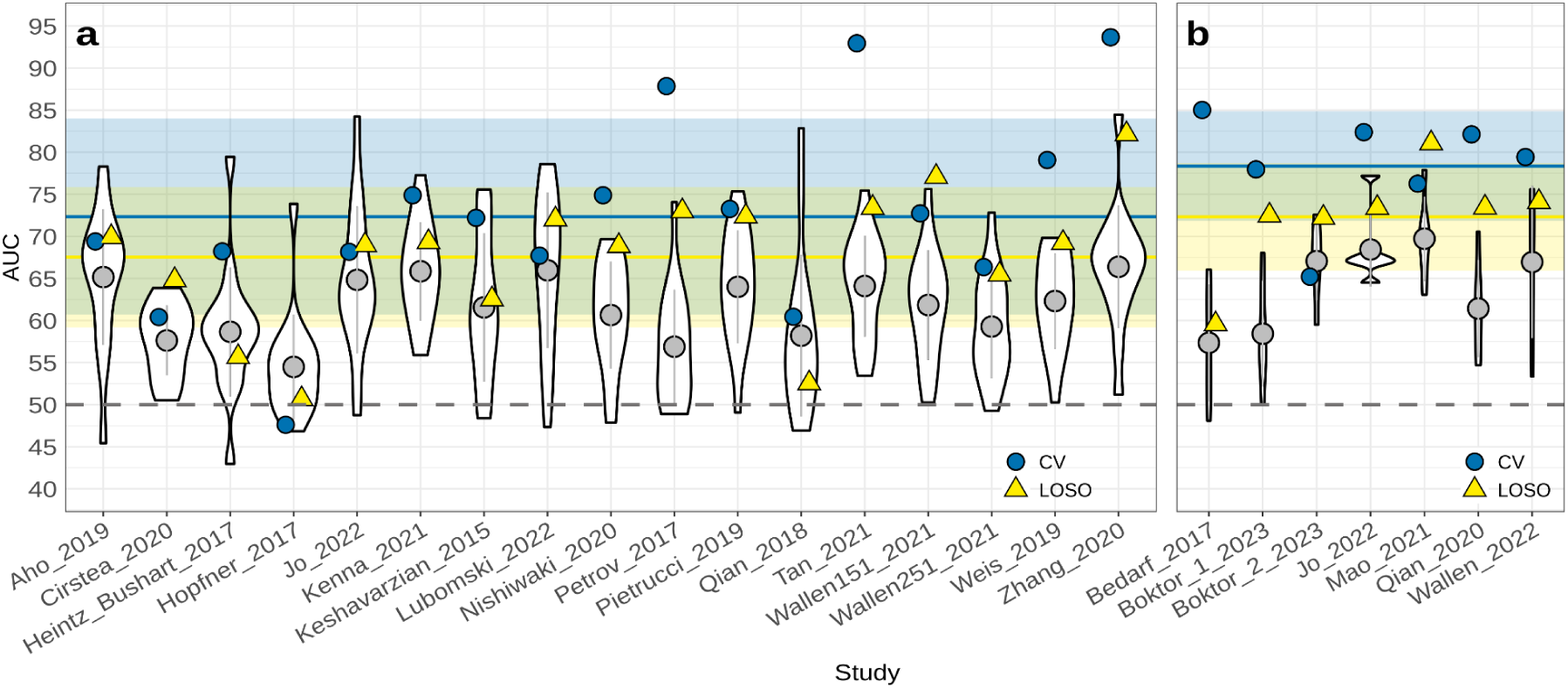
Prediction accuracies of the microbiome-based ML models. Performance of the ML models across datasets for the 16S (a) and SMG (b) data. Violin plots depict the distribution of the ML performances (AUCs) calculated for the study-to-study validation (CSV; where the study-specific models were tested on data from all other studies). Within the violin plots the average AUC and the standard deviation are reported (grey dots and lines). Blue circles and yellow triangles indicate the AUC for the within-study cross-validation (CV) and the leave-one-study-out validation (LOSO), respectively. LOSO AUCs are obtained by training ML models on all but one dataset, which is then used to evaluate the model as a hold-out set (study on the X-axis). The horizontal shaded areas and lines indicate the standard deviation and average of the AUCs for LOSO (yellow) and CV (blue). The grey dashed line marks the 50% AUC threshold indicating random guessing.

Finally, we examined whether by pooling data across studies model performance could be improved in comparison to models trained on single studies. To do this we performed a leave-one-study-out (LOSO) validation, in which all data are combined with the exception of data from one study that is then used to evaluate the model as a hold-out dataset. For both 16S and SMG data the LOSO model performances were significantly higher than those obtained through CSV (Fig 2; Welch two sample t-test: 16S t = -2.8, df = 17.8, p-value = 0.01, metaG t = -3, df = 8.9, p-value = 0.01). For both data types, LOSO AUCs were not correlated with the number of samples used for model training (Pearson correlation, 16S: t = -1.5, df = 15, p-value = 0.17; SMG: t = -0.37, df = 5, p-value = 0.73). Between data types, average LOSO AUCs for SMG were higher than those obtained for the 16S data (Welch two samples t-test: t = -1.5, df = 14.7, p-value = 0.15; LOSO AUC average: SMG = 72.3 ± 6.39, 16S = 67.5 ± 8.35). The results of the LOSO validation lead us to hypothesise that models built for data collected within the same population (i.e. continent) might be more similar to each other, meaning they might share more discriminatory features with each other compared to models built on data from different continents. If true, these differences might influence the CSV and LOSO performances. To verify this hypothesis we extracted the feature coefficients from all Ridge models built on the 16 and SMG data and used them to build an ordination based on Canberra distances (Fig S8). We then tested whether the continent of origin was able to significantly explain the clustering of the samples using a Permanova analysis. Although the datasets partially clustered according to the continent of origin, this was statistically significant only for SMG data (Permanova p-value: 16S = 0.40; SMG = 0.04), for which however only a few datasets are currently available .

**Fig 3.**
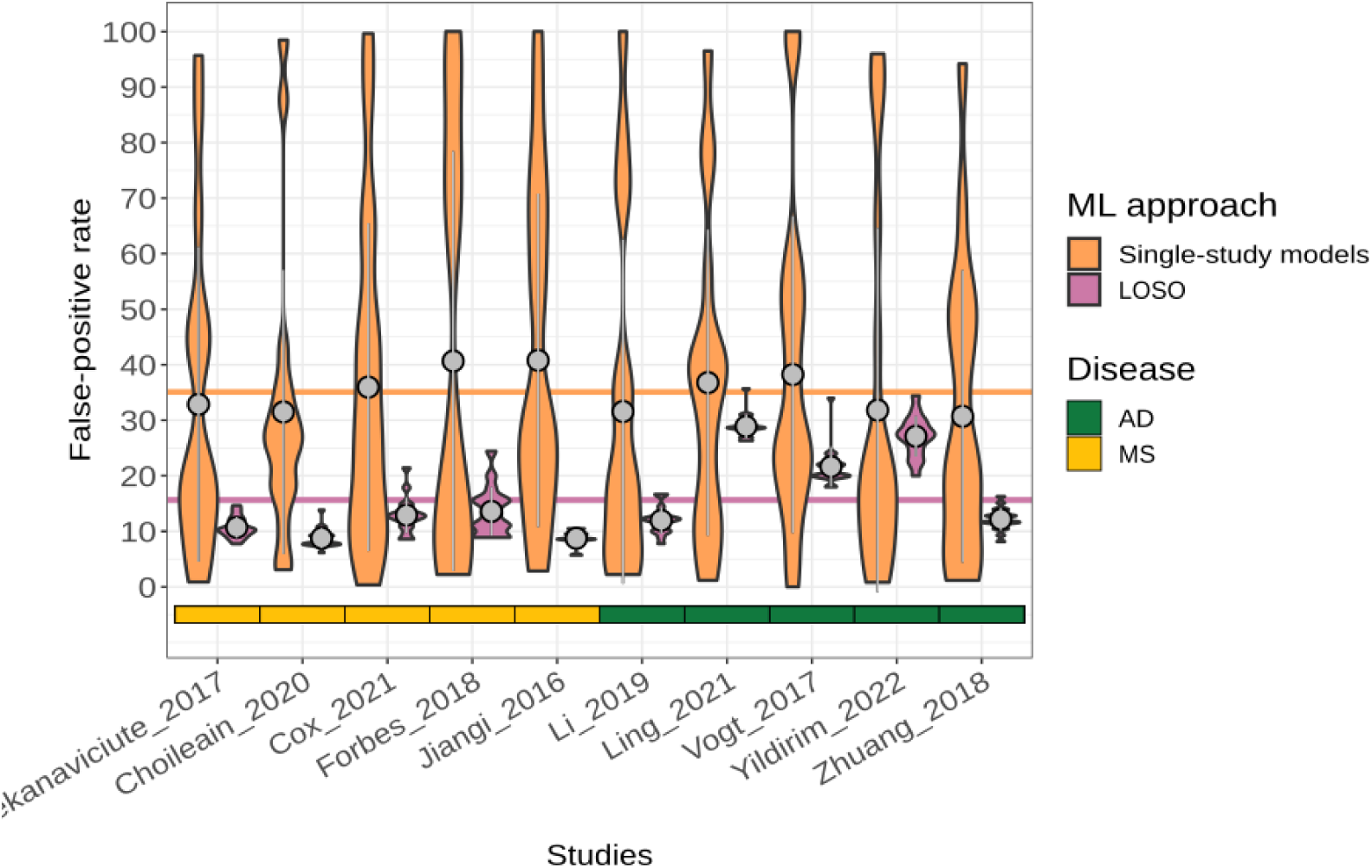
Disease specificity of classification models is significantly improved by pooling data from multiple studies. Violin plots depict with density curves the proportions of samples spuriously predicted to come from PD patients in datasets of other neurodegenerative diseases (to assess the disease specificity of the respective PD classification models). All classification models (Ridge regression) trained on individual PD studies as well as on pooled data (LOSO) were evaluated for false-positive prediction (FPR) rates on datasets obtained for other diseases (AD = Alzheimer’s disease; MS = multiple sclerosis). Models were originally adjusted to a 10% FPR on PD datasets and an FPR above this level in patients with other diseases indicates lack of disease specificity as previously established by Wirbel et al13. Within the violin plots the average and standard deviation are reported (grey dots and lines). On average, LOSO models were found to be much more disease specific (average FPR = 15.7%, horizontal purple line) than models trained on a single PD dataset (average FPR = 35.1%, horizontal orange line).

### Cross-disease prediction

Our results show that some ML models can accurately discriminate PD patients from controls, but models are generally study-specific and do not perform well when tested on external datasets. Additionally, it is of relevance to assess if model predictions are specific for PD, that is to check to what extent they wrongly predict patients affected by other neurodegenerative diseases than PD. To assess cross-disease prediction rates, we tested each PD model on data obtained from studies about other neurodegenerative diseases. We performed this cross-disease validation using only 16S data due to limited availability of SMG data. We tested all Ridge models built for each PD 16S dataset on data from Alzheimer’s disease (AD) and Multiple Sclerosis (MS). The observed cross-prediction rates (assessed in terms of the false positive rate, FPR, on AD and MS samples and compared to the PD-internal FPR of 10%) varied greatly across the PD-specific ML models, ranging from 0% to almost 100%, with an average of 35.1% (Fig 3). However, cross-disease prediction drastically improved when the LOSO models were used, as the average FPR was reduced to 15.7%, which is only moderately higher than the expected 10% FPR for PD-internal control groups. Our finding that disease specificity of ML models can be significantly improved by training on data pooled across multiple studies confirms earlier reports on the effectiveness of this approach^37^.

**Fig 4.**
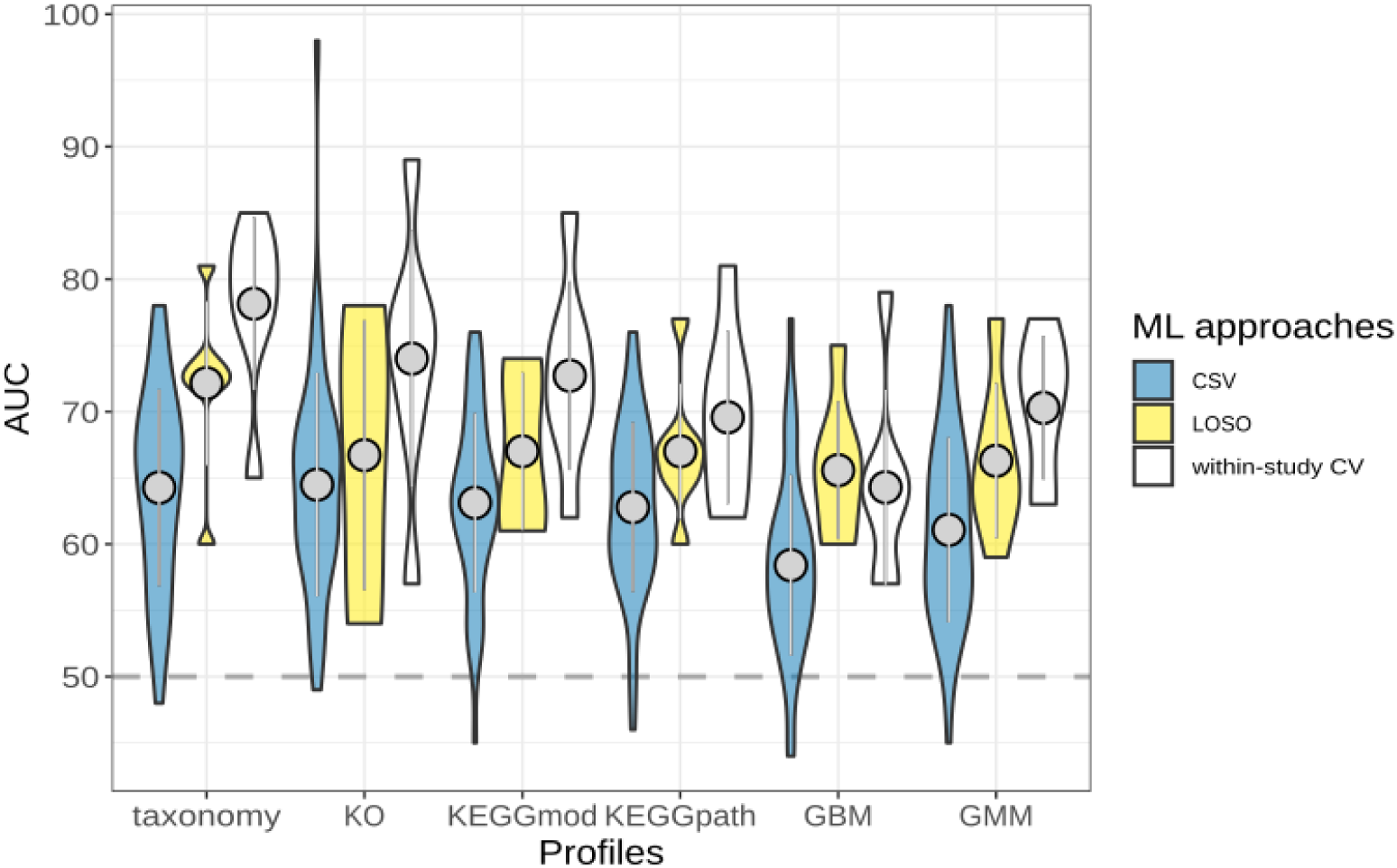
Taxonomic profiles perform better than functional profiles in discriminating PD from controls. Violin plots depict with density curves the AUC values obtained across ML approaches (within-study cross-validation, CV; study-to-study validation, CSV; and leave-one-study-out validation, LOSO). Within the violin plots the average AUC and the standard deviation (grey dots and lines) are reported. Performances for ML models (Ridge regression classifiers) built on taxonomic (mOTUs) and functional profiles are reported. The grey dashed line marks the 50% AUC threshold indicating random guessing. KO = KEGG orthologous gene families; KEGGmod = KEGG modules; KEGGpath = KEGG pathways; GMM = gut metabolic modules; GBM = gut-brain axis modules; taxonomy = mOTUs_v3 profiles.

### Comparison between taxonomic and functional microbiome profiles

Not only taxonomic profiles, but also functional microbiome profiles, which are derived from SMG data to capture a broad range of metabolic and other pathways, have previously been used for building classification models. It remains a disputed question, whether models based on functional profiles are more accurate than those obtained from taxonomic profiles^13,17^. In our meta-analysis, we found that models built on taxonomic profiles perform in general better than those built on functional profiles (Fig 4). This finding was consistent across ML models built on different types of functional profiles, namely KEGG orthologous groups (KOs), KEGG modules, KEGG pathways, gut metabolic modules (GMM), or gut-brain axis modules (GBM) (Fig 4, Fig S9-S13). The latter are manually curated microbial metabolic pathways shown to play important roles in gut health and the gut-brain axis^39,40^. Among the different functional profiles, KOs performed slightly better in discriminating PD from controls, and had slightly higher CSV performances compared to KEGG modules and pathways, GMM, or GBM (Fig 4, Fig S9-S13). As for the taxonomic profiles, CSV accuracy values were overall considerably lower than those obtained for the within-studies CV (Fig 4, S9-S13). Overall these results indicate that the use of functional profiles does not improve the classification accuracies or the ML model portability (as assessed by CSV) when compared to taxonomic profiles.

### Taxa associated with PD

To identify taxa consistently associated with PD across datasets, we performed a meta-analysis on relative abundances of gut microbial taxa (Fig 5; Fig S14, S15, Supplementary data 1 & 2). This was done by calculating Generalised Odds Ratios (Gen. Odds) and pooling the effect sizes using random effect meta-analysis. We then corrected *p-values* with the false-discovery rate (fdr) and selected a *q-value* cutoff of 5%. In addition, we used the datasets with available metadata to verify using linear models whether the microbiome features associated with PD might be potentially confounded by sex, age, or medication usage. In both 16S and SMG data, taxa within the *Lachnospiraceae* family belonging to the genera *Roseburia*, *Blautia*, and *Fusicatenibacter* were strongly depleted in the microbiome of PD patients. Similarly, we detected the genus *Agathobacter*, within the *Lachnospiraceae* family, to have strongly reduced abundance in the 16S datasets. Affiliated to this genus are mOTUs 03657 and 12366 which were depleted in PD patients across the SMG datasets. Similarly, *Faecalibacterium* (family *Ruminococcaceae*) was found strongly and consistently depleted in PD. The high-resolution profiling we performed for the SMG data allowed us to identify multiple species within the *Faecalibacterium* genus and multiple strains within the *Faecalibacterium prausnitzii* species (mOTUs 06112, 06110, 06109, 06108) depleted in the microbiome of PD patients. The species showing the strongest depletion in PD patients belonged to the *Butyricicoccus* genus (Fig 5, Supplementary data 2). However, this association was not detectable in 16S datasets, even though the corresponding family *Butyricicoccaceae* was reported as PD depleted in our previous meta-analysis^12^. Finally, we found the genus *Prevotella* and the species *Prevotella copri* (mOTUS 03701) to have reduced relative abundance in PD patients, whereas the species *Prevotella corporis* (mOTUs 02390) was enriched in PD.

**Fig 5.**
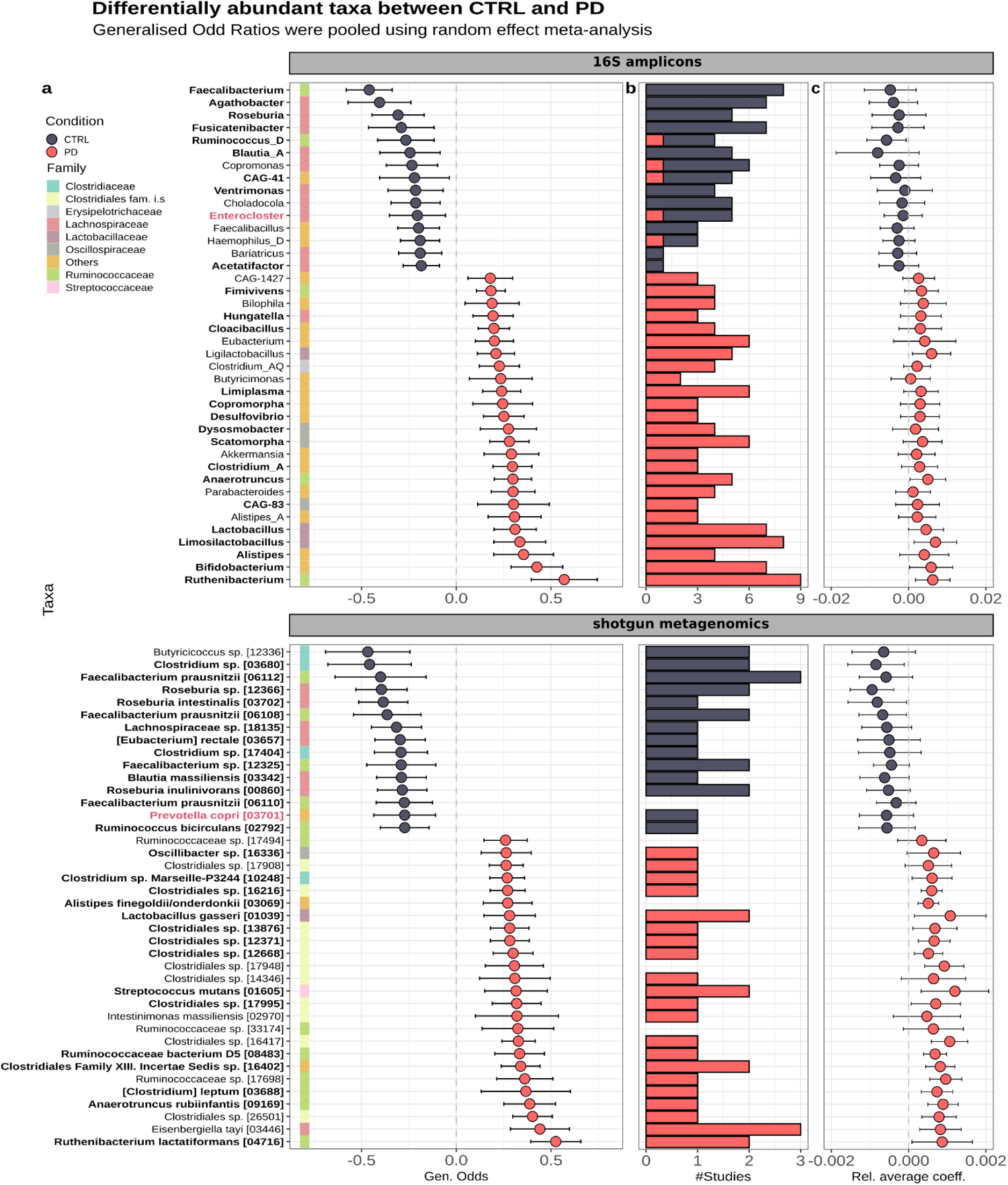
Taxa showing significant differences in abundance between PD and controls (CTRL). Univariate testing was performed independently for each dataset using Agresti Generalised Odd ratios on the taxa obtained from 16S and SMG profiles. Results were pooled using random effect meta-analysis and displayed together with 95% confidence intervals (a). (b) Number of studies in which each taxon significantly differs in abundance between PD and CTLR. (c) Average Ridge coefficients and standard deviations calculated from all models built on each dataset. Only 40 taxa with the largest effect sizes are shown. Genera and mOTUs with similar direction of enrichment are displayed in bold. The genera Enterocloster and Prevotella have one related mOTU enriched in PD and one in CTRL and are thus reported in red.

Good concordance between 16S and SMG data was also observed for the taxa enriched in PD. For example, the 5 genera with the strongest enrichment in the 16S datasets had related mOTUs enriched in the SMG data (Fig 5, 6; Fig S14, S15, Supplementary data 1 & 2). Differently from previous studies^12,15,17,23^, we detected in both 16S and SMG data the *Ruthenibacterium* genus and the *Ruthenibacterium lactatiformans* species as the most enriched taxa in the PD gut microbiome. Similarly, taxa within the genera *Alistipes*, *Anaerotruncus*, *Enterococcus*, *Porphyromonas*, *Scatomorpha*, *Limiplasma*, *Bifidobacterium, Christensenella*, *Streptococcus* were all consistently enriched in the 16S and the SMG datasets. Several differences in the taxa associated with PD were also observed between the two sequencing methods (Supplementary data 1 & 2). For example, we detected the potential pathogenic species *Turicibacter sanguinis* (mOTU 04703), and multiple species within the order Clostridiales enriched in PD samples, but the respective genera did not have a significantly increased abundance in the 16S datasets. As we used an up-to-date version of the Genome Taxonomy Database (GTDB v207) to taxonomically classify the 16S-derived ASVs, we could discriminate with higher resolution, compared to previous studies, the genera within the *Lactobacillus* sensu-lato that are more abundant in PD patients. The genus *Lactobacillus* has been recently reassessed taxonomically and several new genera have been created^41^. Among these, *Limosilactobacillus*, *Lactobacillus*, *Lacticaseibacillus*, and *Ligilactobacillus* were all enriched in the gut microbiome of PD patients (Fig 5, S14, Supplementary data 1).

We observed that taxa differentially abundant in the pooled meta-analysis had significant abundance shifts in only a fraction of the individual datasets, in agreement with previous findings^12,16^(Fig 5b, Supplementary data 1 & 2). This is most likely due to both the variability observed across studies and the low statistical power derived by the small sample size of many datasets we re-analysed. Only a minor fraction of the taxa associated with PD (<22%) resulted potentially confounded by sex, age, or medication usage, and in general the taxa with the strongest abundance shift were not affected by covariates (Supplementary Data 10-13). When comparing the results of differential abundance tests applied to each taxon with their influence in the classification models (i.e. their coefficient size in the Ridge regression classifiers), we found these to be remarkably consistent. The sign of model coefficients for these taxa mostly matched the direction of association from univariate analysis although variability in the average Ridge weights across datasets were evident (Fig 5c, Fig S14c, S15c, Supplementary data 1 & 2).

**Fig 6.**
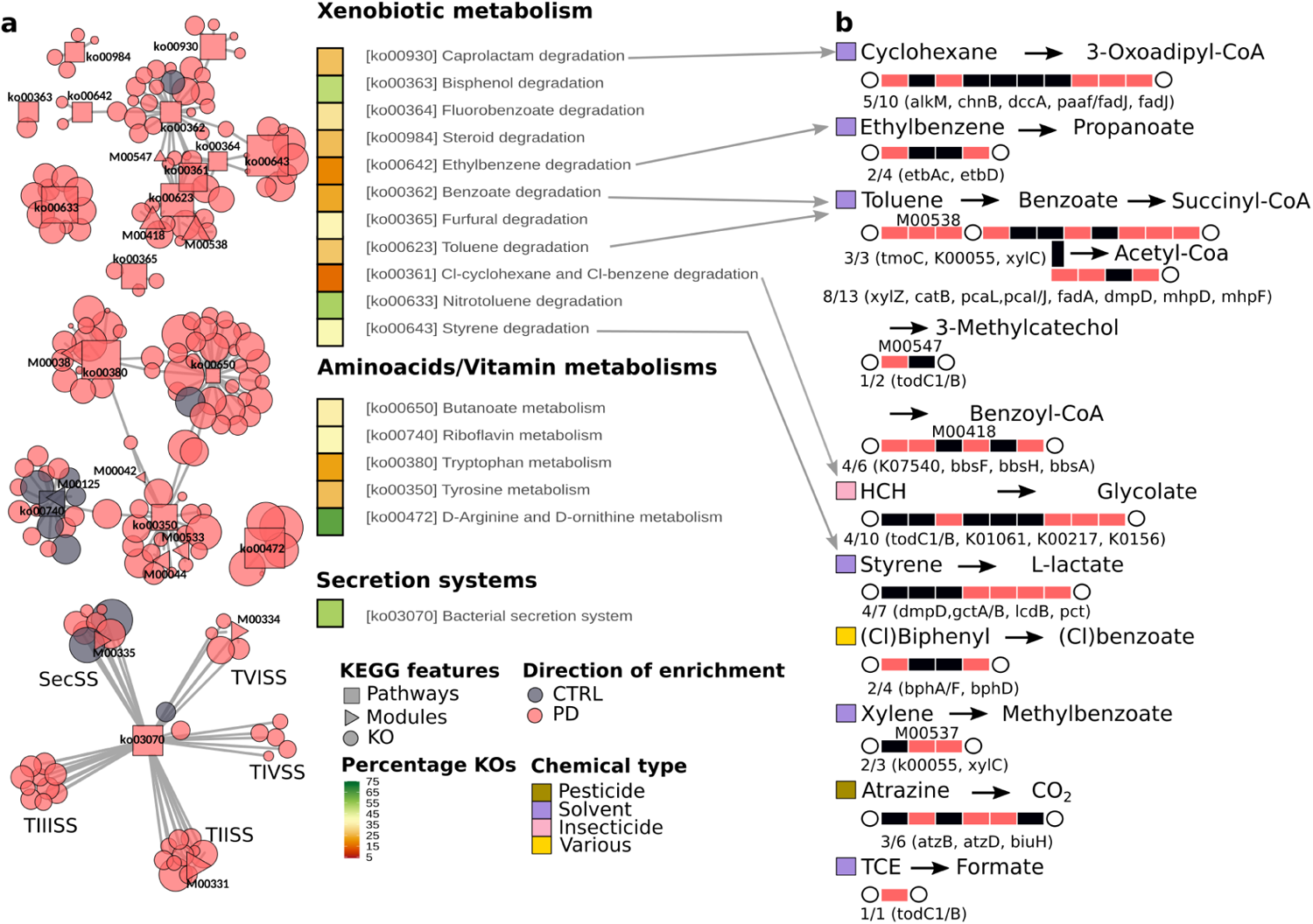
KEGG functionalities with significant differences in abundance between PD and controls (CTRL). Selected KEGG-based functionalities are displayed using a network in which the relationships between pathways, modules, and KO are indicated (a). The type of functionalities and the direction of enrichment is displayed using shapes and colours, respectively (see key). The size of each node in the network is proportional to the effect size (larger effect size = stronger enrichment). Only pathways and module nodes are labelled. Heatmaps indicate the proportion of differentially abundant KOs, out of all KOs in the pathway. TIISS = type II secretion system; TIIISS = type III secretion system; TIVSS = type IV secretion system; TVISS = type VI secretion system; SecSS = Sec secretion system. For the xenobiotic metabolisms, we report here a summary of the most representative reactions catalysed by KOs enriched in PD (b). Each rectangle corresponds to an enzyme or a complex of enzymes, representing a metabolic reaction within the respective KEGG pathway. Rectangles are depicted in red when at least one of the KOs catalysing the enzymatic reaction has been detected as enriched in PD. The number of reactions with KOs enriched in PD and the respective KO gene names are reported below each graph. Circles depict metabolites and xenobiotics, which are colour-coded to reflect their use by humans.

### Gut microbial gene functions associated with PD

To explore changes in gut microbial functionalities in PD patients relative to controls, we extended the differential abundance meta-analysis, performed as described above, to microbial genes and pathways as defined by the KEGG orthologous groups (KO), KEGG modules, KEGG pathways, GMMs, and GBMs (Fig 6, Fig S16-S18, Supplementary data 3-7). We complemented this approach with an enrichment analysis to detect those pathways that were significantly enriched in KOs showing differential abundances between PD and controls (Supplementary data 8). Below we will highlight those gut microbial functions with a possible relation to Parkinson’s aetiology or symptomatology.

Modules related to the degradation of complex polysaccharides and sugars were strongly depleted in PD (KEGG modules: M00631, M00061, M00081; GMM: MF0001, MF0003, MF0004, MF0022, MF0010, MF0018, MF0002; KEGG pathways enriched in Kos depleted in PD: ko00040, ko00520; Supplementary Data 4, 6-8). These findings are consistent with previous reports^15,18^ and align well with the depletion of bacteria able to ferment complex carbohydrates into SCFAs in PD. In contrast to these results, some functionalities related to the production of propionate and butyrate were enriched in PD (MF0093, MF0094, MF0089, Supplementary table 6). However, low levels of SCFAs have been observed in faeces of PD patients^42^, thus future studies aimed at integrating metagenomics and metabolomics are required to elucidate how the microbiome shapes the SCFAs concentrations in the gut of PD patients.

Several pathways involved in the metabolism of amino acids had a significant difference in abundance between PD and controls (Fig 6a; Fig S16-S18; Supplementary data 3-8). These pathways are particularly relevant in the context of the gut-brain axis, as amino acids, especially the aromatic ones such as tryptophan and tyrosine, are precursors of neurotransmitters that have altered concentration in PD^43^. Moreover, animal experiments have shown that concentrations of neurotransmitters in the host gut and brain can be affected by the gut microbiome through the modification of amino acid pools^44–46^. Our results suggest that the PD gut microbiome has an increased ability to degrade tryptophan as genes encoding enzymes involved in this process were significantly enriched in PD gut metagenomes, while those involved in tryptophan synthesis were depleted (KEGG pathway: ko00380, KEGG Module: M00038, GBM: MGB049, MGB004, MGB005 and GMM MF0025; Fig S17-S18; Supplementary data 3 and 7). Our data hint towards an increase in microbial tyrosine turnover in the gut of PD patients, as we detected a significantly higher abundance of genes for both tyrosine degradation and synthesis (KEGG pathway: ko00350, KEGG modules: M00044, M00042, and M00040, and the GMM MF0027; Fig S17-S18; Supplementary data 3-7). Within this pathway, the gene coding for tyrosine decarboxylases TyrDC (K22330) was enriched in PD gut metagenomes. This enzyme catalyses the degradation of the main PD medication, L-dopa, in *Lactobacillus* sp. and *Enterococcus* sp.^47^, suggesting that PD medication regimes might influence the metabolism of the PD gut microbiome. Indeed, our analysis suggested that some of these pathways might be affected by PD medications, but not L-dopa, as well as by other medication usage (Supplementary data 10-13, Fig. S19). Significant alterations in abundances were also observed for genes related to the metabolism of glutamine, glutamate, and 4-aminobutyrate (GABA), which are all essential amino acids for brain metabolism and function. PD metagenomes were depleted in genes encoding enzymes for glutamate synthesis and showed a converse enrichment in genes involved in its degradation (enriched in PD: equivalent modules MF0032 and MGB051; depleted in PD: equivalent modules MGB007 and MF0047; KEGG KOs: K01846, K19268, K04835, K00265, K00266, K00284; Fig S17, S18 and Supplementary data 6-7). Finally, enzymes catalysing the degradation of GABA and gamma-Hydroxybutyric, a natural GABA precursor, as well as the last step of the glutamate conversion into succinate through GABA, had significantly higher abundance in PD metagenomes (KEGG module: M00027; KEGG KOs: K00135; GMM MF0076 and GBM MGB039; Fig S15-S16; Supplementary data 3, 6, 7). Although these data might suggest an increased ability of the PD microbiome to degrade GABA, we also detected an enrichment, albeit with smaller abundance shifts, of enzymes catalysing GABA synthesis (GBM: MGB021 and MGB020; KEGG module: M00136) and a potential confounding effect of age on the abundances of these functions (Supplementary Data 13). Together this suggests a complex interplay between production and degradation affecting GABA concentration, and indicates that other omics techniques such as metabolomics are needed to elucidate the dynamics of this metabolite in the gut of PD patients.

PD gut metagenomes were enriched in genes encoding proteins involved in the adhesion to, interaction with, and manipulation of host cells, as well as the resistance against host immune responses. Specifically, the KEGG pathway for bacterial secretion systems was significantly more abundant in PD metagenomes (ko03070; Fig 6a; Fig S15; Supplementary data 5). Secretion systems are complex molecular machineries used by bacteria to release effector proteins in the surrounding environment or into neighbouring cells^48^. Specific types of secretion systems (i.e. types III, IV, VI) are used by pathogens to inject effector molecules into host cells to manipulate their defence and immune systems^49^. Within this pathway, 52.7% of KOs were differentially abundant between PD and controls, with the types II, III, IV, and VI secretion systems showing the clearest enrichment in PD. In agreement with these results, additional KOs related to type VI secretion system and to the *Legionella*-like type IV secretion systems were enriched in PD as well (K11890, K11895-7, K11900, K11909, K12083, K12210-1, K12213, K12217-8, K20531; Supplementary Data 3).

Similarly enriched were several modules and KOs involved in bacterial resistance against cationic antimicrobial peptides (CAMP; KEGG modules: M00730, M00739, M00744, M00723, M00722, M00726, M00725; Supplementary data 4 and 8). CAMPs are important host defence mechanisms produced at sites of infection and/or inflammation^50^. Hence, finding an enrichment of these defence mechanisms suggest an ongoing host immune response towards microbes. Some of the above associations might be potentially confounded by sex and age (Supplementary data 13, Fig S19), with these functions having higher abundances in the older population and in males, which are both known intrinsic risk factors for PD^1,4^. Another way in which bacteria can interact with their host is by producing extracellular structures called curli fibres that are involved in cell adhesion, biofilm formation, and bacterial virulence^51^. Confirming previous findings^15^, KOs for curli fibres showed a significantly higher abundance in PD (K04337-8, K06214, K04334-5; Supplementary Data 3). These amyloid-like bacterial proteins have attracted considerable interest in relation to PD as they have been shown to promote αSyn aggregation and motor impairment in mice^52,53^ suggestive of a causal effect of gut bacteria on PD development. Altogether our results indicate an enrichment of potential pathogenic functions in the gut microbiome of PD patients and suggest an increased activation of host defence mechanisms towards infectious agents.

Finally, our in-depth analyses on gut microbial functions revealed that multiple pathways within the KEGG class *“Xenobiotics biodegradation and metabolism”* were significantly enriched in PD. To the best of our knowledge this has been previously discussed only on the basis of functionalities inferred from 16S data^35^. This is relevant since exposure to environmental xenobiotics (e.g. pesticides, herbicides, solvents) is one of the main non-genetic risk factors for developing PD^4,54–57^. In these enriched pathways, between 15.4 and 52.6% of all pathway-KOs were significantly more abundant in PD (Fig 6a). Some of these KOs (e.g. K04072, K00121, K00170) can be part of the central metabolism and thus might not necessarily take part in xenobiotic degradation. Moreover, it cannot be excluded that these enzymes may metabolise medications taken by PD patients. Indeed we detected a minority of these features to be potentially confounded by medication intake in addition to sex and age (Supplementary Data 10-13, Fig S19). However, we found not-confounded KOs enriched in PD that appear to be specifically involved in the metabolism of environmental xenobiotics: for example the 2-haloacid dehalogenase K01560, which takes part in the degradation of halogenated hydrocarbons (Supplementary data 3). These molecules have been widely used as solvents, industrial chemicals, pesticides, and herbicides, and have been linked with PD before^54,55^. For example, recent epidemiological studies suggested that individuals exposed to water contaminated with trichloroethene (also known as trichloroethylene or TCE) had a 70% increased risk of developing PD^55^. Interestingly, our analysis revealed that PD gut metagenomes were enriched in K03268 and K18089, which encode enzymes that can catalyse the conversion of TCE into formate. Considering the relevance of xenobiotics in PD aetiology, we further inspected the PD-enriched KOs manually to identify those involved in xenobiotic metabolism and related pathways (Fig 6b). In addition to the KOs belonging to PD-enriched pathways, we observed a significant increase in abundance of other KOs involved in xenobiotic metabolism, even though the whole pathway did not pass our significance threshold. For example, PD samples had a higher abundance of the genes *atzB*, *atzD*, and *biuH* (K03382, K03383, K19837; Supplementary Data 3), which encode enzymes that catalyse the degradation of atrazine. Atrazine is a widely used chlorinated pesticide that showed dopaminergic toxicity in rat models^56^. Hence, it is compelling to find a link between the enrichment of microbiome functionalities in PD and known risk factors previously associated with the disease.

## Discussion

In recent years, several studies have suggested that the gut microbiome might be leveraged to support PD diagnosis^17–20,22^. However, a consensus on gut microbiome features associated with PD and a thorough evaluation of microbiome-derived biomarkers for PD is currently missing. The extensive ML validation and optimization we performed here underline that within most study populations the ML models based on the gut microbiome could accurately discriminate PD from controls. However, ML models were study-specific, i.e. poorly generalised to data from other studies (cross-study portability tested using CSV). Cross-study portability was generally higher for the SMG-based models, suggesting that larger technical heterogeneity between 16S studies (i.e. 16S variable region used) might limit comparability. However, it is worth emphasising that PD is a very heterogeneous disease in terms of aetiology, pathophysiology, and symptomatology^6,43,58^ reflected in an ongoing discussion on whether it is appropriate to consider PD a single disease^59,60^. This biological heterogeneity is rarely accounted for in microbiome studies, as samples come from patients i) affected by different PD-types; ii) having different PD severity resulting in different lifestyles; iii) having individual medication regimes; iv) reporting disparate histories of medical conditions and exposure to risk factors (e.g. xenobiotics). All these aspects might exert heterogeneous influences on microbiome composition, and contribute to the low study-to-study portability and high variability in accuracy of ML models across datasets we observed. Moreover, these differences can potentially confound the associations between microbiome features and disease conditions. To thoroughly assess potential confounders, the scarcity of standardised publicly available metadata poses a severe limitation. Here we did analyse the metadata available for some studies, which suggested that a minor part of PD associated microbiome features may be confounded (<28.2%). It is worth noting that to identify all potential confounding effects larger datasets with standardised metadata are required. Hence, caution is needed in concluding on the value of gut microbiome biomarkers for PD diagnostics. On a positive note, the data from Bedarf et al.^18^ that comprises L-dopa-naive early PD patients, allowed us to build ML models that classify PD cases with high accuracy. In their study, the authors ruled out an overall influence of PD medication on the microbiome abundances. Hence, it is reasonable to expect that the gut microbiome of these patients more closely resembles the one of undiagnosed/early PD patients. Therefore, the results related to this particular study suggests that microbiome signatures may capture truely PD-associated signals. However, for diagnostic/predictive purposes the potential of the gut microbiome as disease biomarker would need to be explored in larger multi-center studies of drug-naive early PD patients, or of high-risk individuals, as has been recently attempted in two independent investigations^61,62^. Another important prerequisite for future clinical application of microbiome-derived biomarkers for PD is their disease specificity, i.e. their capability to distinguish the PD microbiome signature from those of other neurological diseases. Towards this aim, we demonstrate that pooling data from multiple studies allows for building PD-specific ML models that have low rates of false-positive predictions when tested on microbiome profiles from other neurodegenerative diseases.

Our large scale meta-analysis further contributed to a better understanding of PD processes to which the gut microbiome may contribute. First, we found the gut microbiome of PD patients to be depleted in bacteria known to ferment complex carbohydrates into SCFAs and in pathways involved in complex carbohydrates degradation. While this is in agreement with earlier studies^12,15,16,18,36^, we report this depletion in the largest and most diverse dataset so far analysed, which strongly suggests that it is a common feature in PD across patient populations. Low levels of SCFAs have been linked to compromised gut health, increased permeability and inflammation, as well as prolonged transit time and have been often recorded in faeces of PD patients, who often suffer from compromised gut health^9,42,63,64^. However, the fact that some functions related to SCFAs production were enriched in PD indicates that caution needs to be used in concluding metabolic capabilities of the microbiome based only on metagenomic data. Future multi-omics approaches integrating e.g. metagenomic, metaproteomics, and metabolomics are required to clearly establish how the gut microbiome contributes to metabolite concentrations in the gut of PD patients. Our data indicate that another general feature of the PD gut microbiome is the enrichment of lactic-acid producing bacteria (e.g. *Lactobacillus* sensu-lato, *Bifidobacteirum,* and *Ruthenibacterium*), in agreement with earlier findings^12^. Some *Lactobacillus* strains encode the enzyme tyrosine decarboxylases TyrDC (K22330) which catalyse the degradation of the main PD medication L-dopa^47^. Hence, it might be speculated that the increased abundance of these bacteria is a consequence of their ability to use L-dopa. Here, we verified at a larger scale that among the lactic acid-producing bacteria enriched in PD, TyrDC is encoded only by some taxa within the genus *Lactobacillus* sensu-lato and *Bifidobacterium* (Supplementary Data S9). Therefore, the use of L-dopa alone appears insufficient to explain the enrichment of these bacteria in PD, as their abundances were also not associated with PD medications. Whereas lactic acid-producing bacteria are generally considered beneficial commensals, some of them have also been found enriched in other inflammatory conditions affecting the gut (i.e. IBD) and it has been suggested that they might take advantage of microbiome imbalance in a proinflammatory environment^65,66^. Our finding of their increased abundance in PD further adds to the open question of whether lactic acid-producing bacteria may under some circumstances play a pathological role and contribute to deterioration of gut health in specific diseases.

In PD gut metagenomes we detected an enrichment of type III, IV, VI secretion systems, which are a hallmark of pathogenic bacteria. Our finding aligns well with previous studies reporting an enrichment of potentially pathogenic bacteria in the PD gut microbiome^15,23^, which we partially replicated here. Secretion systems are used by pathogenic bacteria to, amongst others, modulate the host immune response during infection and can cause an activation of inflammatory response and an increase in gut permeability^67,68^. As a first line of defence in the innate immune response against infections, the host can produce CAMP, which are broad spectrum antimicrobials also involved in modulating inflammatory responses^50^. Hence, the enrichment of systems used by bacteria to resist CAMP suggests an ongoing response against potential infective agents in the gut of PD patients. This interpretation aligns well with the higher abundance of virulence related genes (i.e. secretion system and curli fibres) we found in PD metagenomes. Infective agents can lead to an increase in gut inflammation and permeability, which are both commonly observed in PD patients^64^. This deterioration in gut health might then contribute to the translocation of proinflammatory signals and cells to the CNS^69,70^. Finding these functions enriched in the PD gut metagenomes across sampling populations is highly relevant as it adds a mechanistic perspective on how the gut microbiome might contribute to the deterioration of gut health and favour the spread of pathogenic processes along the gut-brain axis. Recently, the connection between gut microbiota, gut health and CNS has emerged as an important aspect affecting neurodegeneration and ageing. For example, faecal microbiota transplantation between aged and young mice showed that the aged donor microbiota increases gut permeability, systemic inflammation, and accelerates age-associated CNS inflammation in young mice^71^. Interpreting our results in the light of these recent experimental findings we can speculate that the gut microbiome of PD patients has an increased pathogenicity potential, which could trigger a pro-inflammatory response and compromise the integrity of the gut epithelial barrier. The compromised gut health and integrity can then facilitate the translocation across the gut epithelium of toxic compounds such as toxic chemicals (see below) and bacterial proteins such as curli fibres. Toxic molecules and bacterial proteins can then more easily reach the CNS stimulating αSyn aggregation, Lewy’s body formation, neuronal toxicity, and neuroinflammation.

Strikingly, the extensive functional metagenomic analyses we performed here revealed many microbial pathways and enzymes involved in xenobiotics degradation to be enriched in PD metagenomes. Although some enriched genes, such as *xylC*, *todC1*, *todB* (KOs: K00141, K03268, K18089) are found exclusively in KEGG xenobiotic metabolism, they can be involved in the degradation of multiple molecules (Toluene, Nitrotoluene, Xylene, Ɣ-Hexachlorocyclohexane, TCE). Hence, the enrichment we observed in these functions does not allow us to pinpoint the specific xenobiotic types that might have contributed to selecting these signatures. However, the enrichment of pathways involved in xenobiotic degradation we observed suggests that the PD microbiome has been exposed to and has adapted to these chemicals. Although we cannot exclude that the enrichment of these pathways is a microbiome adaptation to the medications taken by PD patients, our findings align well with current epidemiological data, which indicates that exposure to such environmental xenobiotics is an important risk factor for developing PD^4,54–57^. There are several conceivable ways in which the observed alterations in gut microbial xenometabolism may be an adaptation to and/or actively modulate environmental exposures. On the one hand, the composition of the gut microbiome might be directly altered as a consequence of exposure to these chemicals^72,73^. In agreement with this first hypothesis, recent experimental data showed that rats exposed to TCE showed signs of PD pathology^55^ and a concomitant gut microbiome enriched in *Bifidobacterium* and a depleted in *Blautia*^57^, similar to the microbiome changes we observed here in human PD patients. On the other hand, it is an intriguing question if or to which extent gut microbial metabolization alters the toxic effects on dopaminergic neurons and the neuroinflammation which some of these chemicals induce^56,57^. Are gut bacteria producing toxic chemicals during the catabolism of these xenobiotics? Besides hypothesising a potential detoxification ability of the gut microbiome, it is not unlikely that some catabolites may have instead an increased toxicity, as has been reported for industrial chemicals and food dyes^72,73^. Since the gut microbiome is characterised by high inter-individual variability, it might represent a person-specific risk modulator of xenobiotic exposures. This is to say that some people exposed to xenobiotics might have a higher likelihood of developing PD due to specific gut microbial metabolic capabilities resulting in increased neurotoxicity, whereas others may benefit from gut microbial detoxification of environmental chemicals. Further work integrating exposure and microbiome data with experimental work on microbial xenometabolism is warranted to shed light on the likely complex interactions between these two important non-genetic factors. The future integration of these two aspects with human genetic markers may enable personalised risk prediction for the development of PD. In summary, our data provide the most comprehensive overview to date about the taxonomic and functional composition of the gut microbiome in PD and provide future reference for its use as a diagnostic tool.

## Material and Methods

### Selected datasets

We collected 16S amplicon (16S) and shotgun metagenomics (SMG) datasets related to case-control studies that compared the composition of the gut microbiome between PD and controls. We include all studies irrespectively of the inclusion/exclusion criteria used, the typology and severity of PD, and the country of origin. We identified a total of 52 studies from which we excluded all studies that profiled <30 samples, did not make raw data available, or for which it was not possible to assign the samples to patients or controls due to the lack of basic metadata. We managed to match the study of Hopfner et al.^33^ with bioproject PRJEB14928, and included this study in our analyses as well. In total we collected 22 datasets, of which 16 and 6 studies profiled the gut microbiome using 16S and SMG sequencing, respectively. To perform a cross-disease comparison of the ML models built for the 16S data we additionally included datasets related to multiple sclerosis^74–78^ and Alzheimer’s disease^79–83^. We performed this test using only 16S data due to limited availability of SMG data for other neurodegenerative diseases.

### Profiling of the 16S amplicon and the shotgun metagenomic data

All 16S data were analysed using the DADA2 algorithm^84^, yielding amplicon sequence variants (ASVs). Trimming parameters were adjusted for each dataset to meet the different quality of the data. Samples sequenced on different runs were profiled independently to allow a correct estimation of the sequencing error rates. The data from Wallen et al.^23^ were sequenced using two different approaches, one using 150 bp and the other 250 bp reads length. Hence, they were split (Wallen151 and Wallen251) and profiled independently. Taxonomy was assigned using the Naive Bayesian algorithm and the GTDB v_207^85^ database. Finally, data were combined at the genus level while samples with <2000 reads were discarded.

Taxonomy profiling of the SMG data was performed using mOTUs v_3.0^86^. The data from Boktor et al.^16^ contained two independent datasets that were analysed here separately (Boktor_1, Boktor_2). The mOTUs taxonomy was then matched with the GTDB v_207 taxonomy using previously published mapping files (https://github.com/motu-tool/mOTUs/wiki/GTDB-taxonomy-for-the-mOTUs). Data were transformed into relative abundances and data classified as “unassigned” were then removed.

Functional profiling of the shotgun metagenomic data was performed using gffquant v_2.10 (https://github.com/cschu/gff_quantifier) in combination with a reduced version of the GMGC human gut nr95 catalogue^87^ obtained by removing genes that only occurred in less than 0.5% of samples used in building the original human gut catalogue. This reduced the catalogue to 13,788,251 non-redundant genes. Prior to functional profiling, raw reads were cleaned using bbduk v_38.93^88^ as follows: 1) low quality trimming on either side (qtrim=rl, trimq=3), 2) discarding of low quality reads (maq=25), 3) adapter removal (ktrim=r, k=23, mink=11, hdist=1, tpe=true, tbo=true; against the included bbduk adapter library) and 4) length filtering (ml=45). The cleaned reads were screened for host contamination using kraken2 v_2.1.2^89^ against the human hg38 reference genome with ribosomal sequences masked (Silva v_138^90^). The remaining reads were then mapped to the reduced human gut catalogue using BWA-MEM v_0.7.17^91^ with default parameters and name-sorted by samtools v_1.14^92^ *collate*. Alignments were subsequently filtered to >45 bp alignment length and >97% sequence identity. Reads aligning to multiple genes contributed fractional counts towards each matching gene. Alignment counts for a gene were normalised by the gene’s length, then scaled up according to the strategy employed by NGLess (https://ngless.embl.de/Functions.html#count) and propagated to the functional features with which the gene is annotated. The final counts were normalised by dividing against the sum of all mapped reads passing our filtration criteria to obtain relative abundances. For KEGG KOs, we retained only KOs of prokaryotic origin according to KOFAMKoala^93^ prokaryotic HMMs. We additionally filtered both KEGG pathways and modules by retaining those consisting of at least 50% and 60% prokaryotic KOs, respectively.

Gut microbial modules (GMM)^39^ and gut-brain modules (GBM)^40^ were inferred based on KOs via the R package omixerRpm v_0.3.3^94^ using default parameters and a pathway coverage (*minimum.coverage*) of 0.5. We then used the KEGG mapper^95^ portal to map the differentially abundant KOs onto the KEGG pathway maps and verify in which xenobiotic metabolisms they are involved. Finally, we used the protein sequence of the TyrDC enzyme encoded by *Enterococcus faecium* (NCBI ID: QAV53956) to verify whether this enzyme is encoded in the genomes of the lactic-acid producing bacteria enriched in PD. The protein sequence was used to query the NCBI database through *blastp*^96^.

### Statistical analyses

All data analyses were performed in R v_4.2^97^. For both taxonomic and functional profiles relative abundances were used for further analyses. First, both 16S and SMG data were used to build ordinations using Bray-Curtis dissimilarities. Data manipulation and ordination were performed using the phyloseq v_1.40^98^ and vegan v_2.6.4^99^ R packages. Ordinations were built using distance-based redundancy analysis (dbRDA) implemented in the *capscale* function within phyloseq as previously described^12^, with and without conditioning the data by study. Significance of the clustering (for both study of origin and disease condition) was then tested on the Bray-Curtis dissimilarities using permutational multivariate analysis of variance (Permanova, *adonis2* function). Permanova for the disease status was performed by restricting the permutation within datasets.

All differential abundance analyses were conducted on data filtered using the above-described 5% criterion, with the exception of the analyses done for the GMM and GBM for which data were not filtered. Agresti generalised odd ratios (*genodds* v_1.1.2^100^ R package) were used to estimate effect sizes and standard errors in each independent datasets. This statistics, analogous to the U statistic underlying the Mann–Whitney test, is based on ranks and does not make strong assumptions about data distributions. It calculates the odds of the second group having a higher value of the outcome (taxa abundances in our case) than the first group if a pair of observations are randomly selected from a dataset. We used the default settings which split the ties to give odds ratios that are equivalent to the Wilcoxon-Mann-Whitney odds ratios. Estimates were then pooled using random effect meta-analysis (meta v_6.2.1^101^ R package), and *p-values* were then adjusted using *fdr*. Adjusted *p-values* will be referred to as *q-values* in the rest of the manuscript. For the functional data, we additionally performed a gene set enrichment analysis using the generic *enricher* function in the R package clusterProfiler V_4.4^102^. This was used to perform independent hypergeometric tests on the subset of KOs enriched either in PD or in CTRL with the aim of estimating which KEGG pathways were significantly enriched in differentially abundant KOs. Background genes (or universe) were defined as all KO within the KEGG pathways that were represented in our dataset. Enrichment tests were run using *minGSSize=5*, *maxGSSize=500*, *p-values* were adjusted using *fdr*, and alpha was set to 5%.

Due to the sparsity of available metadata we used a subset of datasets to perform a sensitivity analysis and identify microbiome features that might be potentially confounded by other donor features such as age, sex, or medication usage. Although age and sex are risk factors for PD and thus intrinsically associated with the disease^1,4^, we included them in this analysis to account for sampling biases. These analyses were performed for all microbiome features we detected associated with PD in our meta-analyses. To test the effect of medication usage we applied two independent strategies. First, we selected all metadata related to medication usage available from Wallen et al.^15,103^ We then retained only medications used in at least 20% of the participants (11 medications in total) and used them to perform a variable selection using the *regsubsets* function in the leaps v_3.1 R package^104^. This was done for the regressions modelling the abundance of the features as a function of medications and disease status, allowing models with a maximum of 12 variables (including all medications and the disease status). We then selected the variables defining the regressions with the minimum Mallow’s Cp value and used them to build the final linear models (lm_covariates; *feature ∼ medications + PD*; where the term *medications* can include up to the 11 medications we considered). Additional baseline linear models were built for the same dataset including only the disease status (lm_pd; *features ∼ PD*). *P-values* were then corrected using *fdr* and compared between model types (lm_pd vs lm_covariates). All features with a significant association with PD (*q-values* in the lm_pd models < 0.05) which were affected by the correction for medication intake (*q-value* in the lm_covariates ≥ 0.05) were considered as potentially confounded. Second, we selected all metadata on PD medications available for the study of Boktor et al.^16,105^ and build linear mixed models for each medication (*feature ∼ medication + (1 | cohort)*). After correcting *p-values* using *fdr* we selected as potentially confounded all those features that had a statistically significant association with at least one PD medication. While these analyses suggested *some* features to be confounded (Supplementary Data 10-12), we need to note that in particular for PD medication this analysis may not be well-powered to detect *all* confounding effects. For a more thorough confounder analysis, more complete data on medication in PD patients is required. Finally, we tested the confounding effect of sex and age by comparing the significance of the association between microbiome features and PD before (baseline models; *feature ∼ PD + (1|cohort)*) and after accounting for covariates (*feature ∼ sex + age + PD + (1|cohort)*). This analysis was performed for both SMG^15,16,18^ and 16S^21,23–25,27,28,32,34^ datasets with available metadata. Metadata from the study of Bedarf et al.^18^ were obtained from the repository related to the study of Boktor et al.^105^ After correcting *p-values* using *fdr*, we selected as potentially confounded all those features having a significant association with PD in the baseline models (*q-value* < 0.05) which became statistically non significant after accounting for covariates (*q-value* ≥ 0.05). All above analyses have been conducted on log-transformed relative abundances.

### Machine learning approaches

Machine learning models were built using the SIAMCAT v_2.0 pipeline^37^. Model accuracy was assessed using a 10-times repeated 10-fold cross validation. We built models using all available algorithms within SIAMCAT (Ridge, Enet, LASSO, Random Forest, Ridge and LASSO as implemented in LibLinear), and tested performances on data normalised using either log or centred log ratio (clr). In addition we tested the effect of feature filtration on model accuracies by building models using all the above indicated algorithms on datasets filtered to retain only the most commonly detected and prevalent taxa. Specifically, we used datasets filtered by discarding all taxa detected in less than 5%, 10%, 20%, 30% of the samples in 10 and 2 datasets for the 16S and SMG data, respectively. Performance of all ML models was quantified by the area under the receiver operating characteristics curve (AUC). For all ML algorithms tested, study-to-study validation (cross study validation; CSV) was performed by testing the models built on each datasets on every other datasets. Leave one study out (LOSO) validation was performed by combining all but one dataset at the time. The combined data were then used to train Ridge ML models and the left out study was used to test model performances. Differences in AUCs between ML approaches were tested using a two-sample Welch t-test as implemented in the R package stats v_4.2.3^97^. We then tested all study specific and LOSO 16S Ridge models on the 16S data obtained for other neurological diseases. False discovery rates, representing the proportion of samples in the test dataset predicted as PD, were then extracted as previously described by Wirbel et al.^13^. ML models for the functional profiles were built by applying the detection filtration described above at the 5% threshold. For the GMM and the GBM no filtration was applied. ML models for the KOs were built by retaining only 600 features selected based on the AUC criterion within the SIAMCAT workflow for nested supervised feature selection. Finally, for all single datasets Ridge models, we extracted the model’s weights (Ridge coefficients) and divided them by the absolute sum of all feature coefficients to calculate relative weights. Relative weights for each feature were then summarised in the figures by average and standard deviation calculated across datasets. The coefficients were further used to create a non-metric multidimensional scaling (NMDS), based on Canberra distances. The effect of the Continent of origin on the clustering of the samples was tested using Permanova.

To test the effect of sequencing depth on the accuracy of the 16S-based models we additionally rarefied the data at a depth of 2000 using the rtk v_0.2.6.1 R package^106^. ML models were then built and evaluated through the same workflow as described above (both CV and CSV) and compared to models built on non-rarefied data using a paired Welch t-test. Moreover, batch effect removal from the 16S data was performed using the function *adjust_batch* with default parameters in the MMUPHin v_1.10.3 R package^107^ and the function *ba* in the bapred v_1.1. R package^108^ using the methods: *meanceter*, which centres the variables within batches (datasets in our case) to have zero mean; *ratiog*, which divides the variables by the batch-specific geometric mean of the corresponding variable; *ratioa*, which divides variable values by the batch-specific arithmetic mean of the corresponding variable. For each batch correction method we then performed independent CSV and within-study CV using all the ML algorithms reported above. Differences in performance within each algorithm type were tested using a one-way ANOVA, as implemented in the R package stats v_4.2.3^97^. Finally, correlations between AUCs values and the number of samples used to train models for the CV, CSV, and LOSO approaches were tested using the *cor.test* function in the R package stats v_4.2.3^97^.

## Data availability

All data used in the article are either publicly available or have been directly obtained from the authors of the original publications, as specified in Table 1.

## Code availability

The R code used in this manuscript is publicly available on GitHub at https://github.com/StfnRomano/PD_ML_meta.

## Acknowledgements

The authors wish to warmly thank all authors that made the data and metadata available for re-analysis. They are moreover grateful to Michael Zimmermann and members of the Zeller group for inspiring discussions. Additionally, we thank Yan P. Yuan, J. Pečar and the EMBL IT Services HPC for support with high-performance computing. The authors gratefully acknowledge the support of the Biotechnology and Biological Sciences Research Council (BBSRC); this research was funded by the BBSRC Institute Strategic Programme Gut Microbes and Health BB/R012490/1 and its constituent project BBS/E/F/000PR10356. SR was partially funded by an EMBO Scientific Exchange Grant (grant no. 9093). GZ is supported by EMBL, the Federal Ministry of Education and Research (BMBF grant no. 031L0181A), the Deutsche Forschungsgemeinschaft (DFG, German Research Foundation no. 395357507 – SFB 1371). The funding bodies had no role in the study design, execution of the analyses, and data interpretation

## Authors’ contributions

SR conceived the project, conducted bioinformatic and statistical analyses, acquired funding, and drafted the manuscript. JW supported statistical data analyses and contributed to the manuscript. RA supported data analysis, visualisation, and interpretation and contributed to the manuscript. QD performed functional profiling and contributed to the manuscript. CS developed the software to perform functional profiling. AN provided financial support and helped with data interpretation. GZ supervised the work, advised on data analysis, visualisation and interpretation, manuscript drafting, and acquired funding. All authors read and approved the final version of the manuscript.

