## Supplementary_figures for "Machine learning-based meta-analysis reveals gut microbiome alterations associated with Parkinson’s disease"

***Fig S1 | AUC across ML algorithms, normalizations, and taxa filtration thresholds.***
*Results refer to ML models built using 16S amplicon data. Distribution of AUCs is depicted using boxplots, in which 50% of the data (25th to 75th percentile) are within the limits of the box and the thick vertical line within it indicates the data's median (50th percentile).*

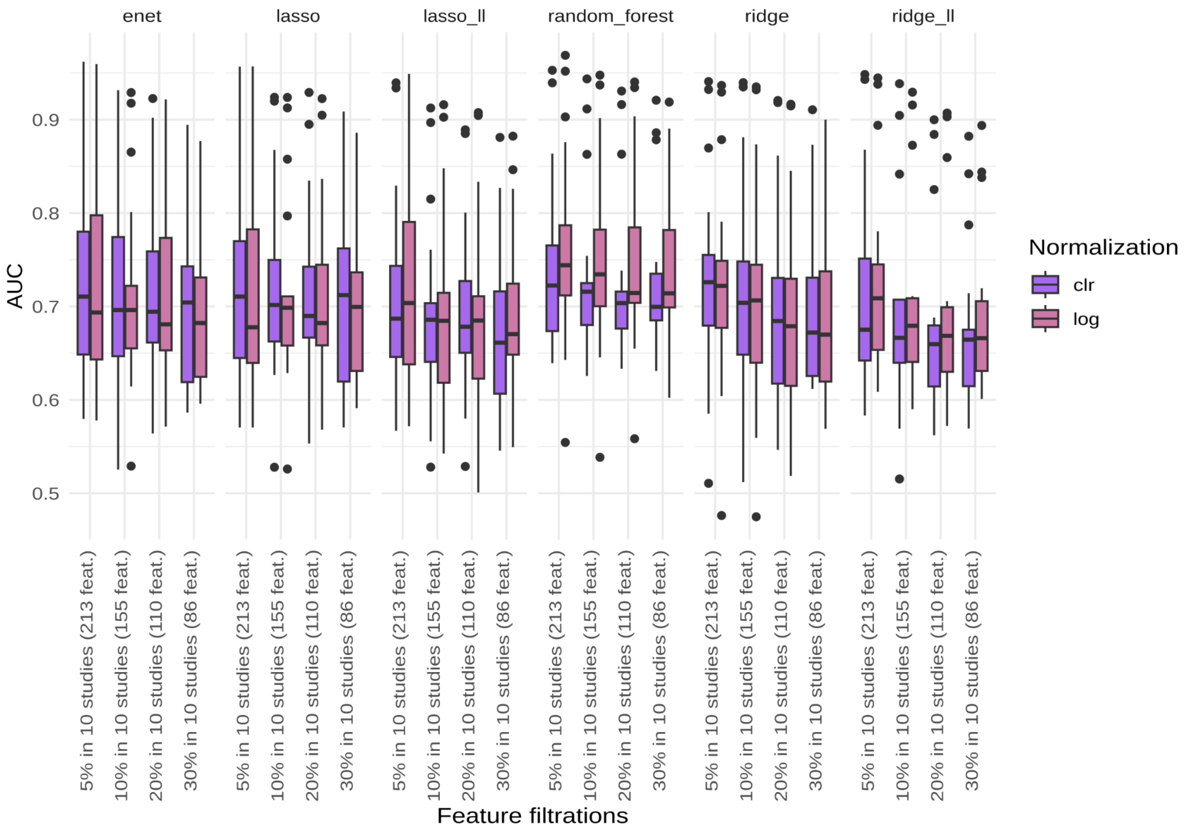

***Fig S2 | AUC across ML algorithms, normalizations, and taxa filtration parameters.*** *Results refer to ML models built using shotgun metagenomics data. Distribution of AUCs is depicted using boxplots, in which 50% of the data (25$^{th}$ to 75$^{th}$ percentile) are within the limits of the box and the thick vertical line within it indicates the data's median (50$^{th}$ percentile).*

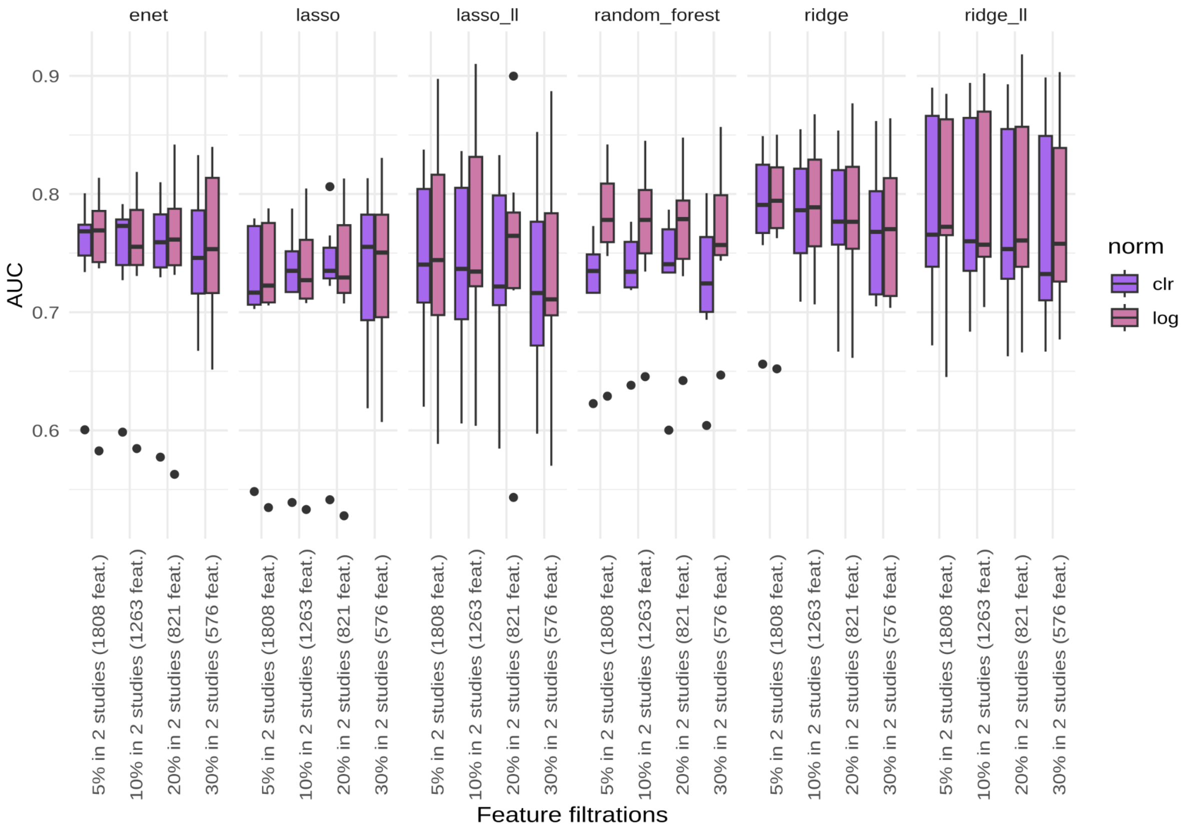

***Fig S3 | AUCs for the study-to-study validation (CSV) performed using 16S amplicon data.*** *Diagonal values indicate the AUCs for the within-study cross-validation (CV). AUCs for the different ML algorithms tested are reported.*

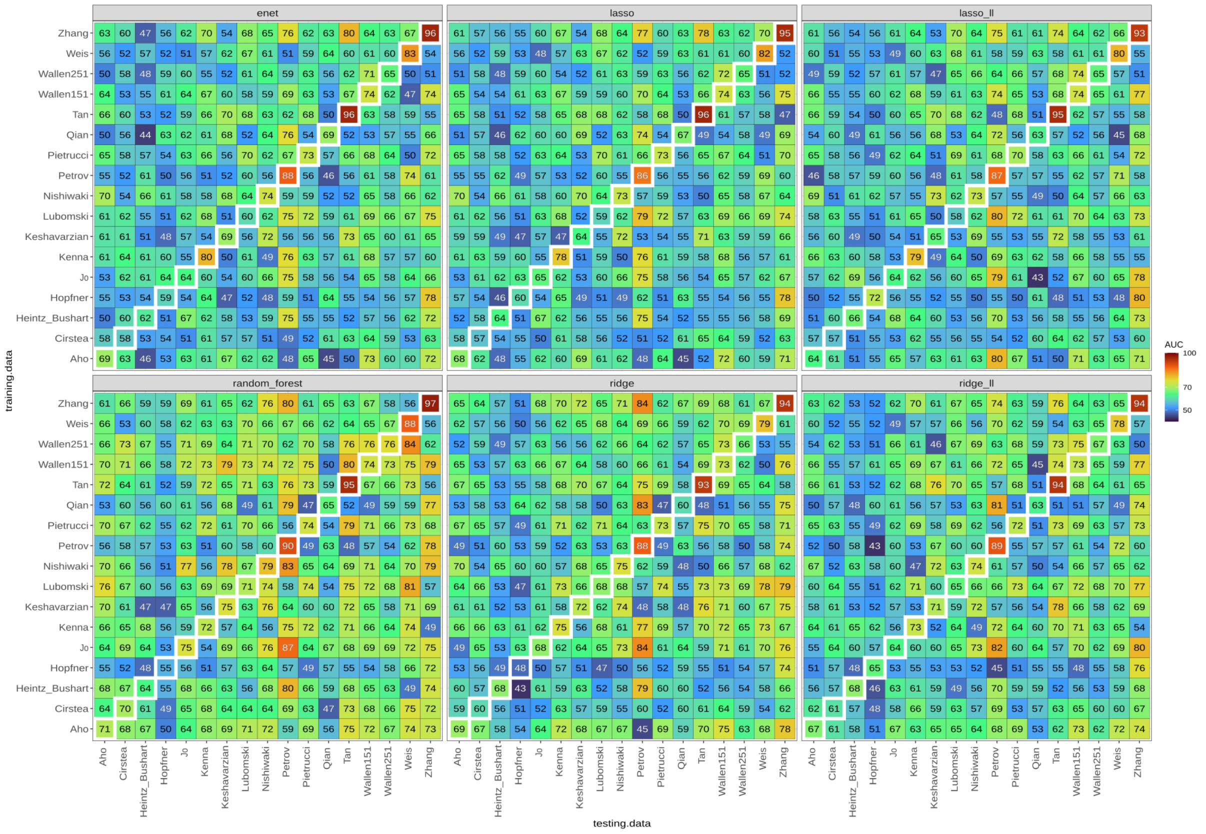

***Fig S4 | AUCs for the study-to-study validation (CSV) performed using shotgun metagenomic data.*** *Diagonal values indicate the AUCs for the within-study cross-validation (CV). AUCs for the different ML algorithms tested are reported.*

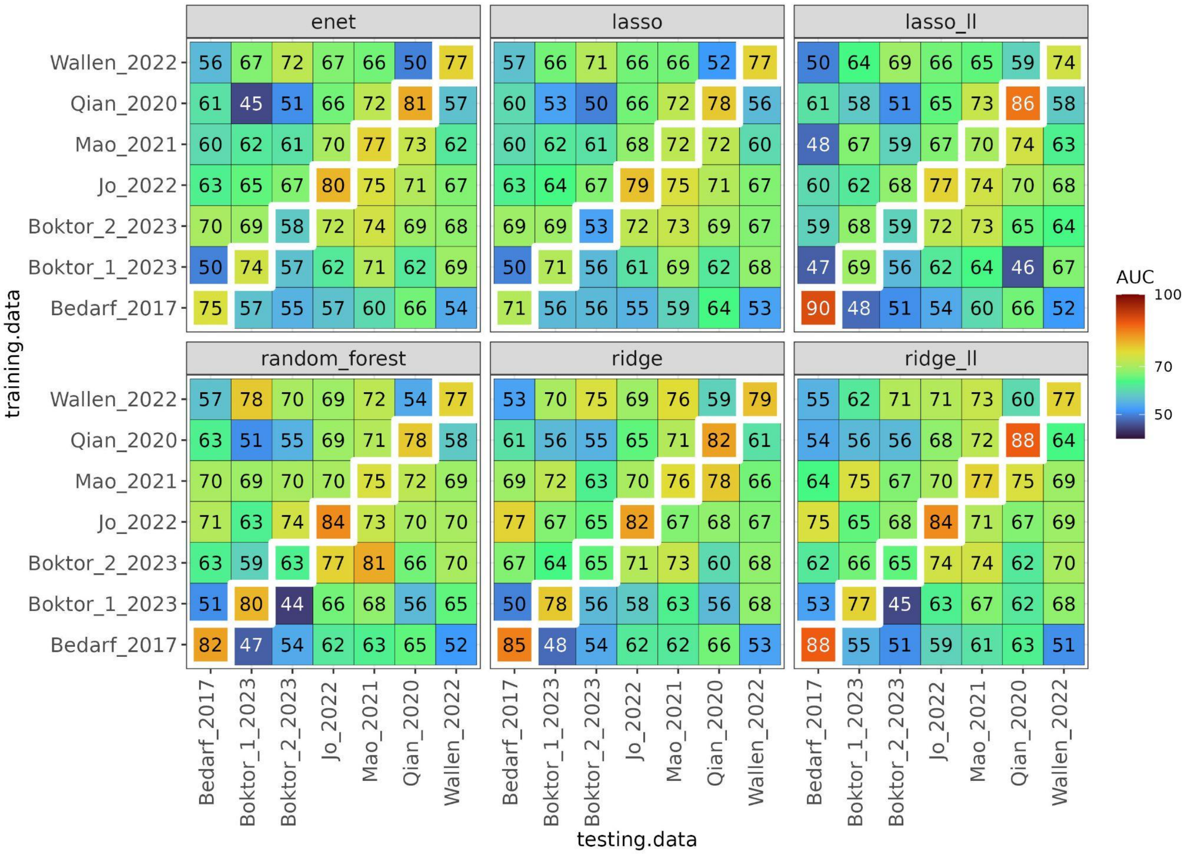

***Fig S5 | Number of reads between conditions and across 16S amplicon datasets.*** *Data distributions are depicted using boxplots, in which 50% of the data (25th to 75th percentile) are within the limits of the box and the thick vertical line within it indicates the data's median (50th percentile).*

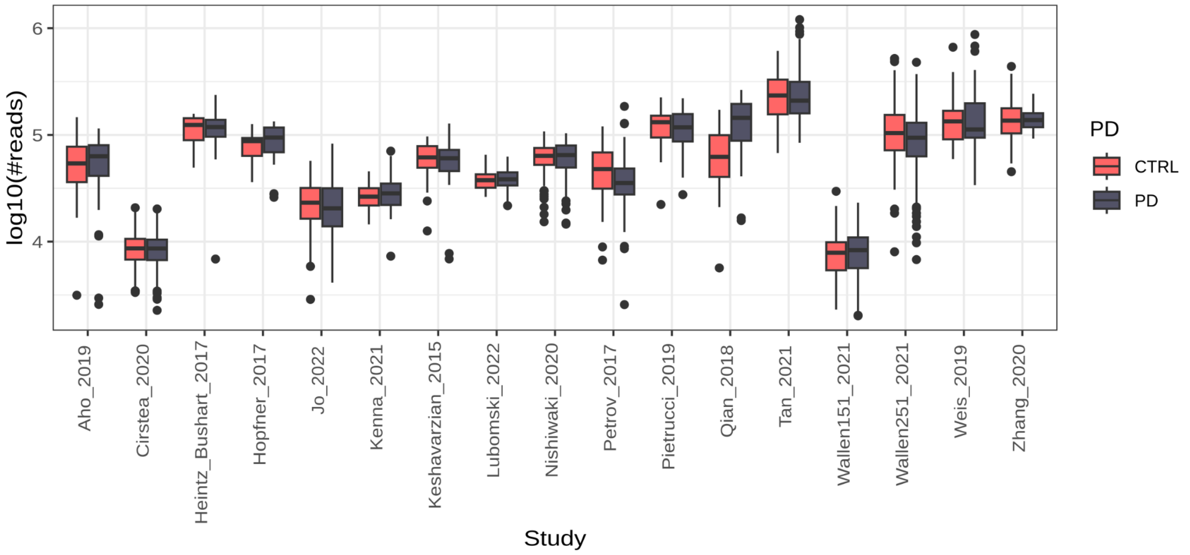


***Fig S6 | Performance comparisons of ML models built on rarefied and not rarefied 16S amplicon data.*** *Data were rarefied to a max depth of 2000, and new ML models were built and used to perform a within-study cross validation (CV) and study-to-study validation (CSV).*

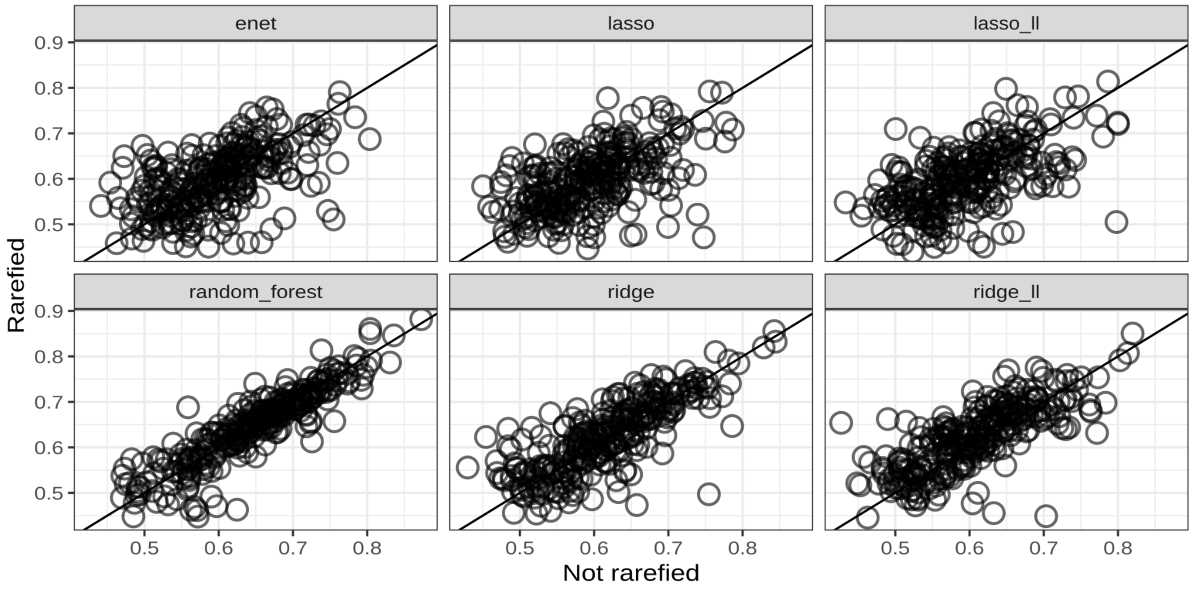

***Fig S7 | Performance comparisons of ML models built on raw and batch-corrected 16S amplicon data.*** *Batch effect was corrected using different label-blind approaches, and the new data were then used to build new ML models used to perform within-study cross validation (CV) a study-to-study validation (CSV). For all algorithms tested, no significant differences were observed between raw data and batch corrected data (ANOVA p-value > 0.79). For details on the different batch correction methods please see the Material and Methods section.*

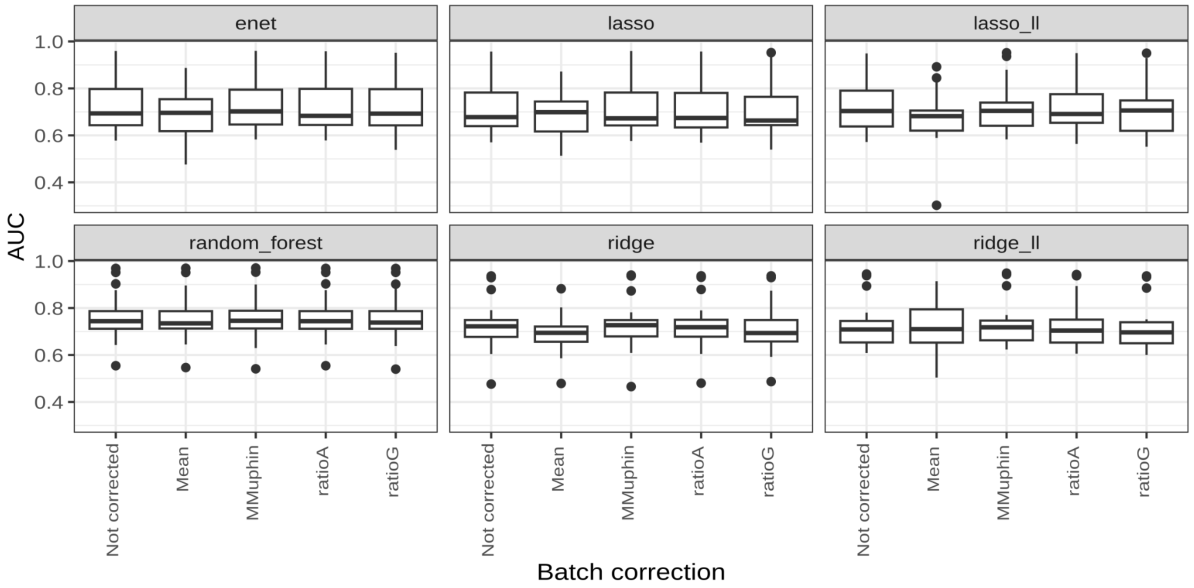


***Fig S8 | Ordination based on Ridge model weights.*** *The ordination referring to the 16S amplicon data is reported in panel A and that referring to shotgun metagenomics data in panel B. Colour of the dots and edges refers to the continent of origin and dataset, respectively. Significance of clustering was assessed using PERMANOVA.*

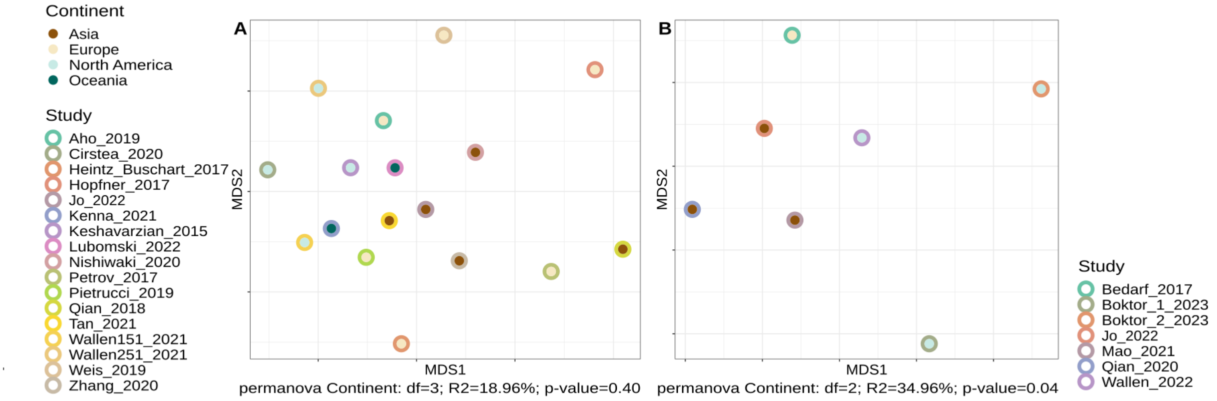

***Fig S9 | AUCs for the study-to-study validation (CSV) performed using KEGG orthologous (KO) inferred from the shotgun metagenomics data.*** *Diagonal values indicate the AUCs for the within-study cross-validation (CV). AUCs for the different ML algorithms tested are reported.*

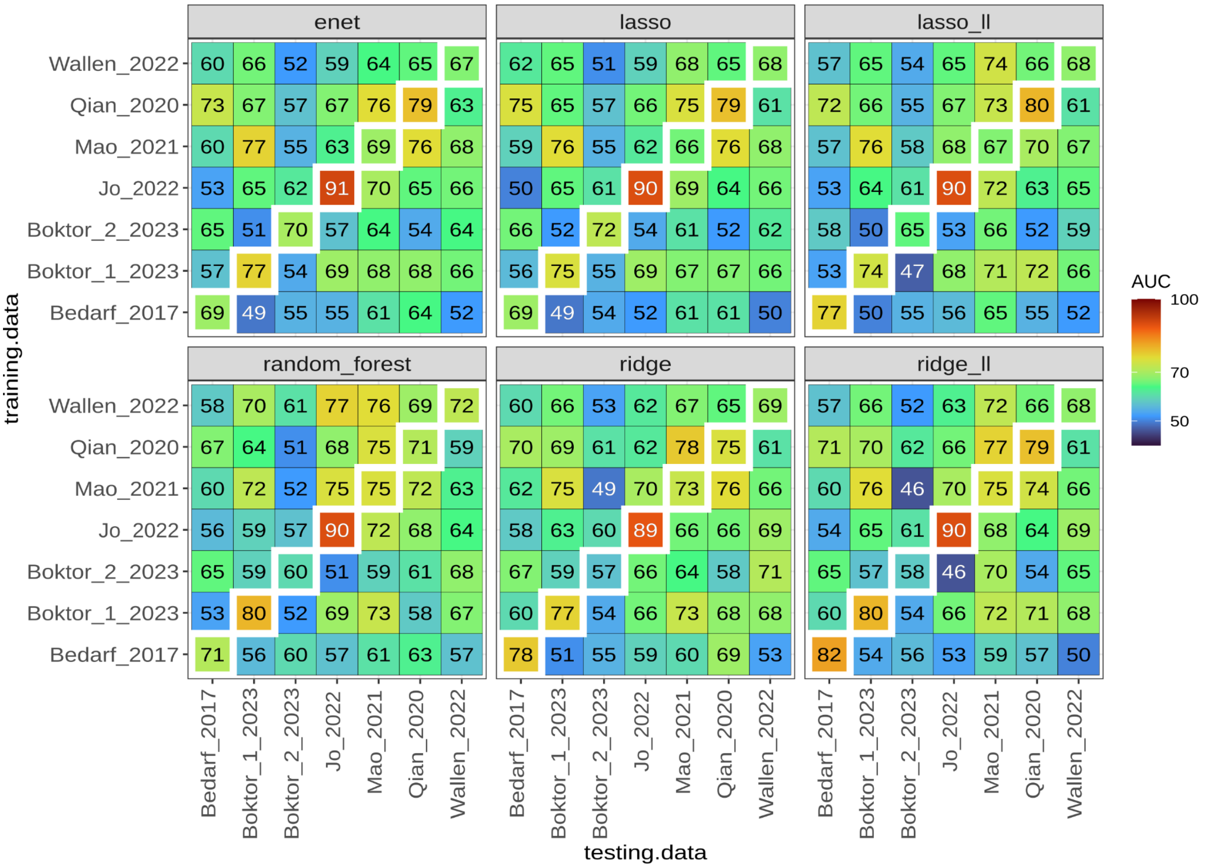

***Fig S10 | AUCs for the study-to-study validation (CSV) performed using KEGG modules inferred from the shotgun metagenomics data.*** *Diagonal values indicate the AUCs for the within-study cross-validation (CV). AUCs for the different ML algorithms tested are reported.*

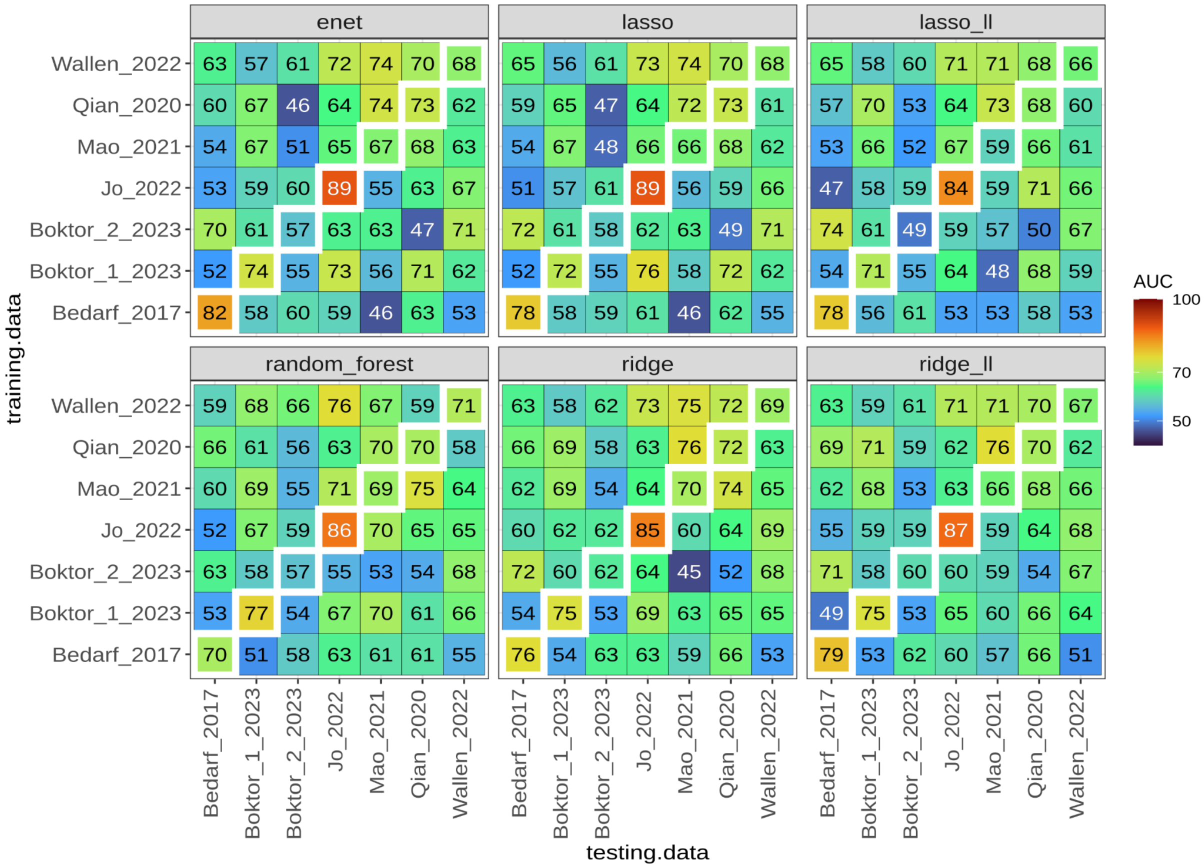

***Fig S11 | AUCs for the study-to-study validation (CSV) performed using KEGG pathways inferred from the shotgun metagenomics data.*** *Diagonal values indicate the AUCs for the within-study cross-validation (CV). AUCs for the different ML algorithms tested are reported.*

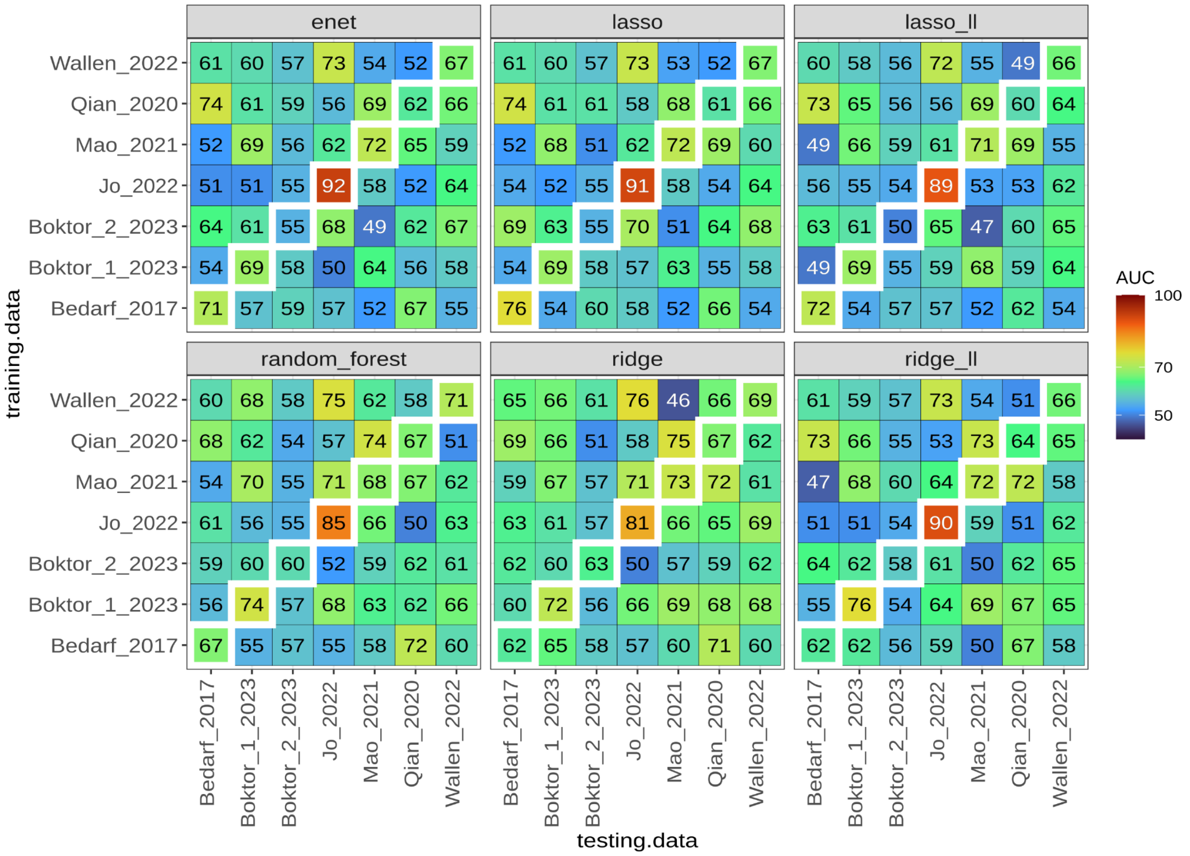

***Fig S12 | AUCs for the study-to-study validation (CSV) performed using gut metabolic modules (GMM) inferred from the shotgun metagenomics data.*** *Diagonal values indicate the AUCs for the within-study cross-validation (CV). AUCs for the different ML algorithms tested are reported.*

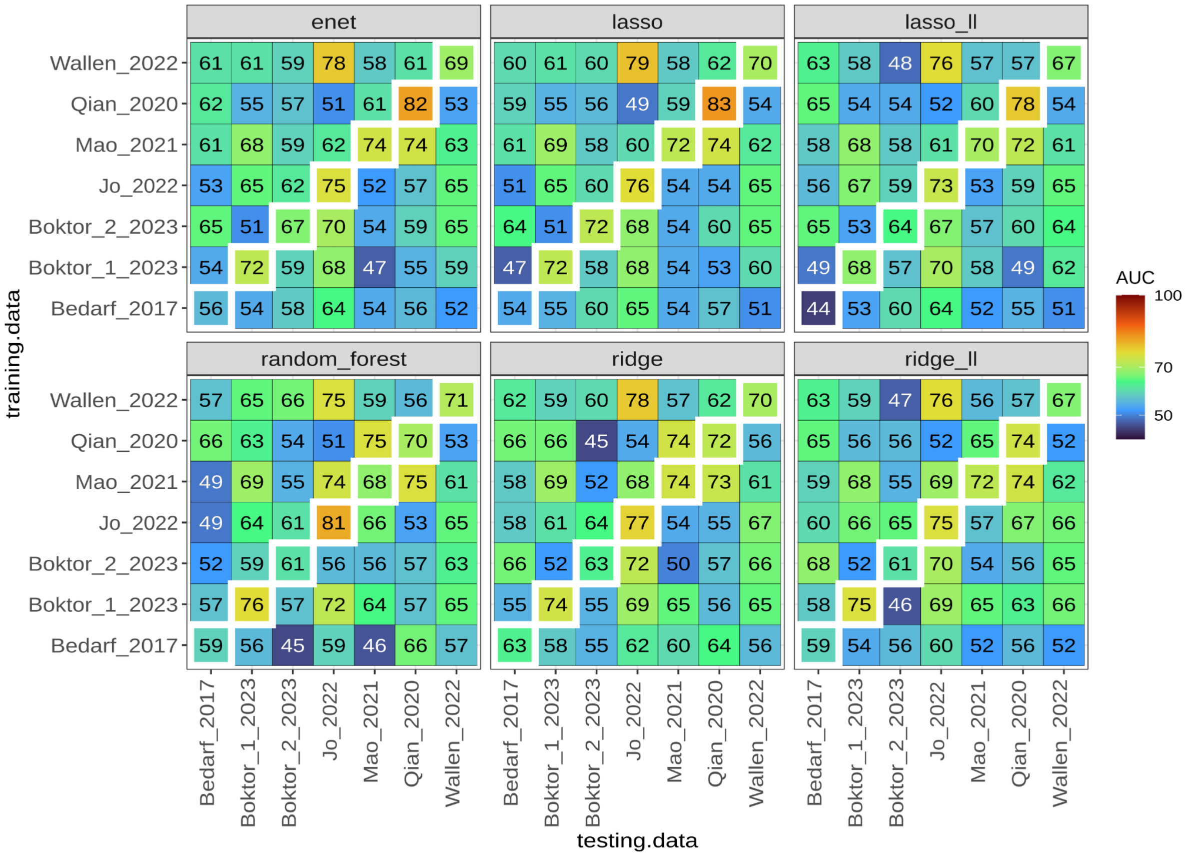

***Fig S13 | AUCs for the study-to-study validation (CSV) performed using gut-brain metabolic modules (GBM) inferred from the shotgun metagenomics data.*** *Diagonal values indicate the AUCs for the within-study cross-validation (CV). AUCs for the different ML algorithms tested are reported.*

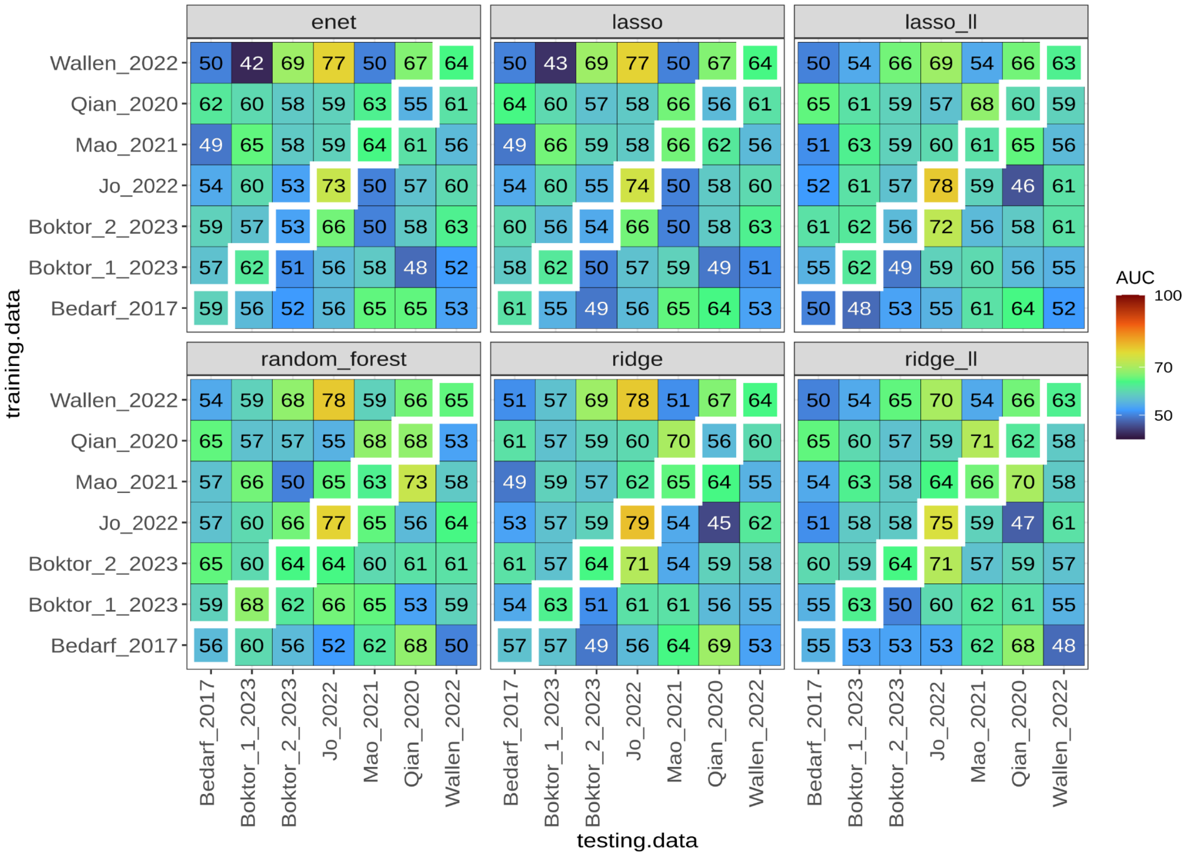

***Fig S14 | Genera showing significant differences in abundance between PD and controls (CTRL).*** *Univariate testing was performed independently for each dataset using Agresti Generalised Odd ratios on the genera obtained from 16S profiles. Results were pooled using random effect meta-analysis and displayed together with 95% confidence intervals (A). (B) Number of studies in which each taxon significantly differs in abundance between PD and CTLR. (C) Average Ridge coefficients and standard deviations calculated from all models built on each dataset. Genera having a similar direction of enrichment as the respective mOTUs (Fig S15) are displayed in bold. The genera Enterocloster and Prevotella have one related mOTU enriched in PD and one in CTRL and are thus reported in red.*

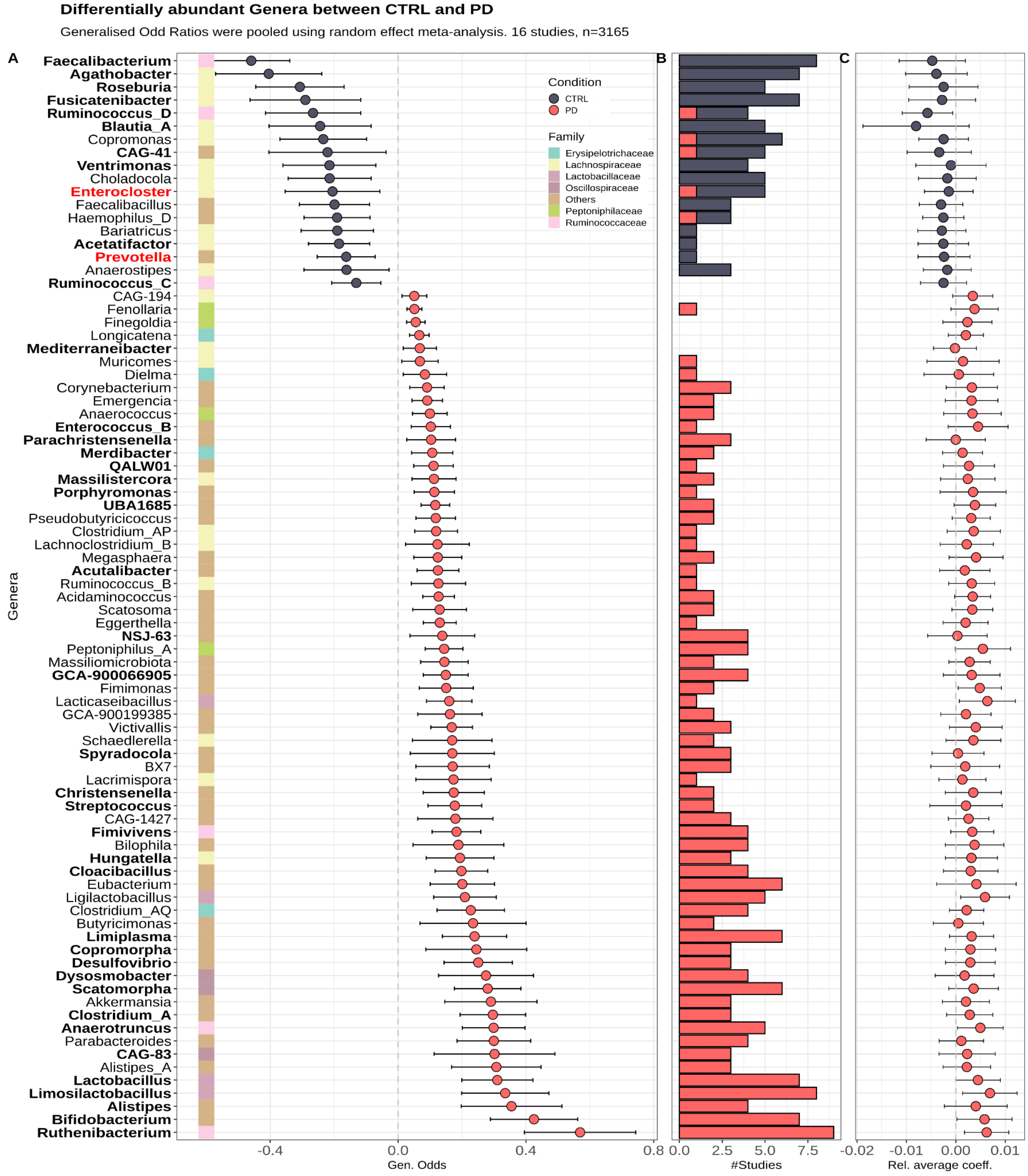

***Fig S15 | mOTUs showing significant differences in abundance between PD and controls (CTRL).*** *Univariate testing was performed independently for each dataset using Agresti Generalised Odd ratios on the mOTUs obtained from SMG profiles. Results were pooled using random effect meta-analysis and displayed together with 95% confidence intervals (A). (B) Number of studies in which each taxon significantly differs in abundance between PD and CTLR. (C) Average Ridge coefficients and standard deviations calculated from all models built on each dataset. mOTUs having a similar direction of enrichment as the respective Genera (Fig S14) are displayed in bold. The genera Enterocloster and Prevotella have one related mOTU enriched in PD and one in CTRL and are thus reported in red.*

**Differentially abundant mOTUs between CTRL and PD**

Generalised Odd Ratios were pooled using random effect meta-analysis. 6 studies, n=1324

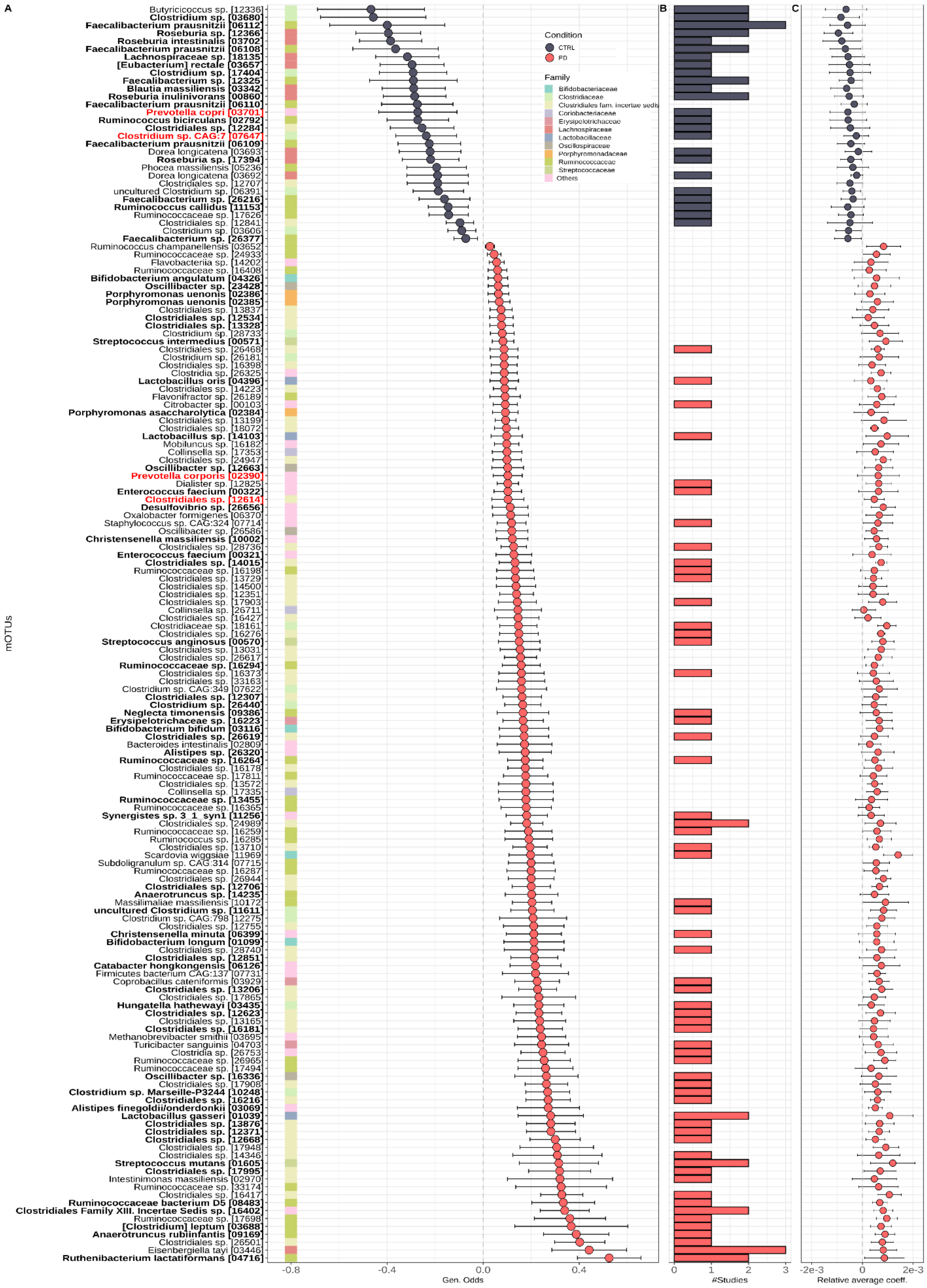

***Fig S16 | KEGG pathways showing significant differences in abundance between PD and controls (CTRL).*** *Univariate testing was performed independently for each dataset using Agresti Generalised Odd ratios on the KEGG pathways inferred from SMG profiles. Results were pooled using random effect meta-analysis and displayed together with 95% confidence intervals (A). (B) Number of studies in which each pathway significantly differs in abundance between PD and CTLR. (C) Average Ridge coefficients and standard deviations calculated from all models built on each dataset.*

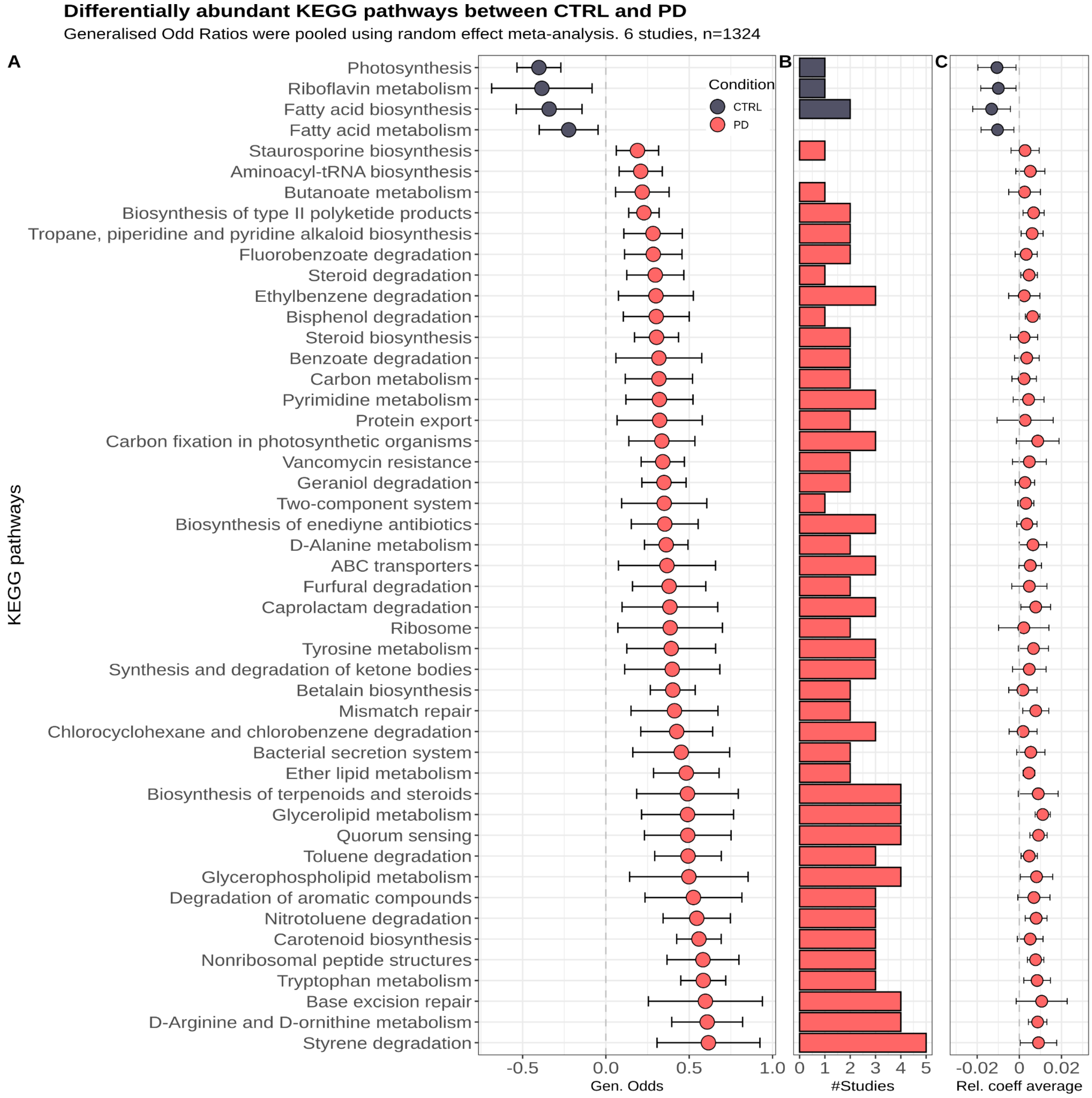

***Fig S17 | GMM showing significant differences in abundance between PD and controls (CTRL).*** *Univariate testing was performed independently for each dataset using Agresti Generalised Odd ratios on the GMM inferred from SMG profiles. Results were pooled using random effect meta-analysis and displayed together with 95% confidence intervals (A). (B) Number of studies in which each module significantly differs in abundance between PD and CTLR. (C) Average Ridge coefficients and standard deviations calculated from all models built on each dataset.*

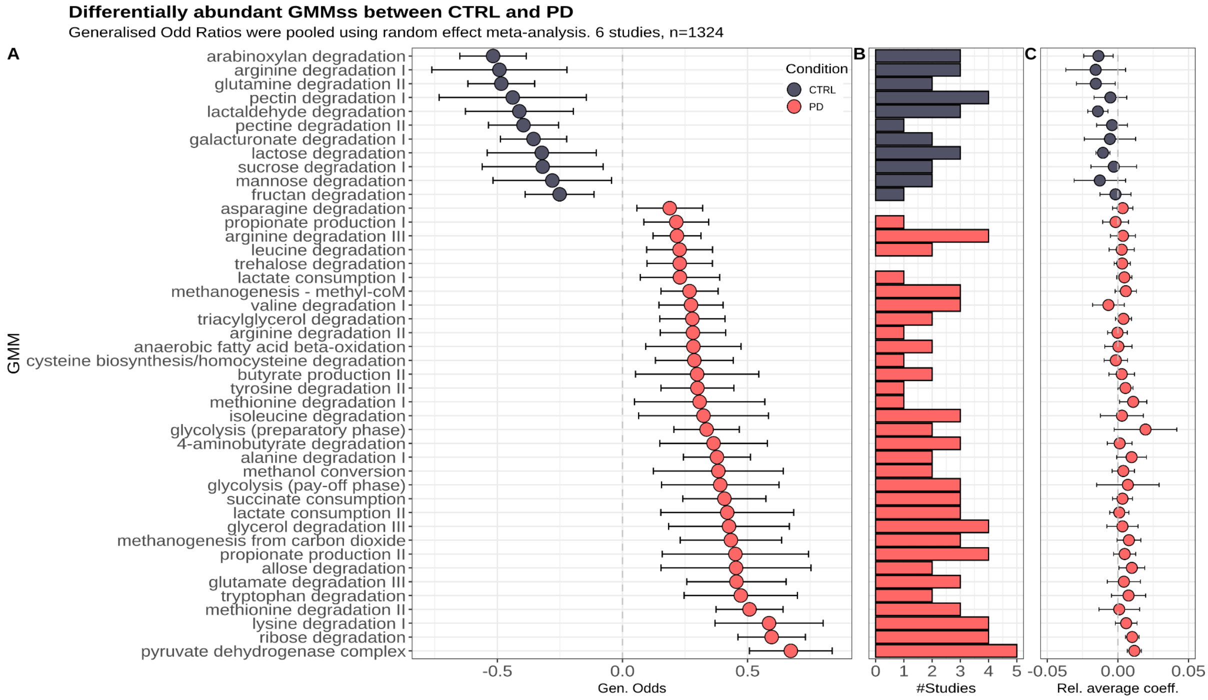

***Fig S18 | GBM showing significant differences in abundance between PD and controls (CTRL).*** *Univariate testing was performed independently for each dataset using Agresti Generalised Odd ratios on the GBM inferred from SMG profiles. Results were pooled using random effect meta-analysis and displayed together with 95% confidence intervals (A). (B) Number of studies in which each module significantly differs in abundance between PD and CTLR. (C) Average Ridge coefficients and standard deviations calculated from all models built on each dataset.*

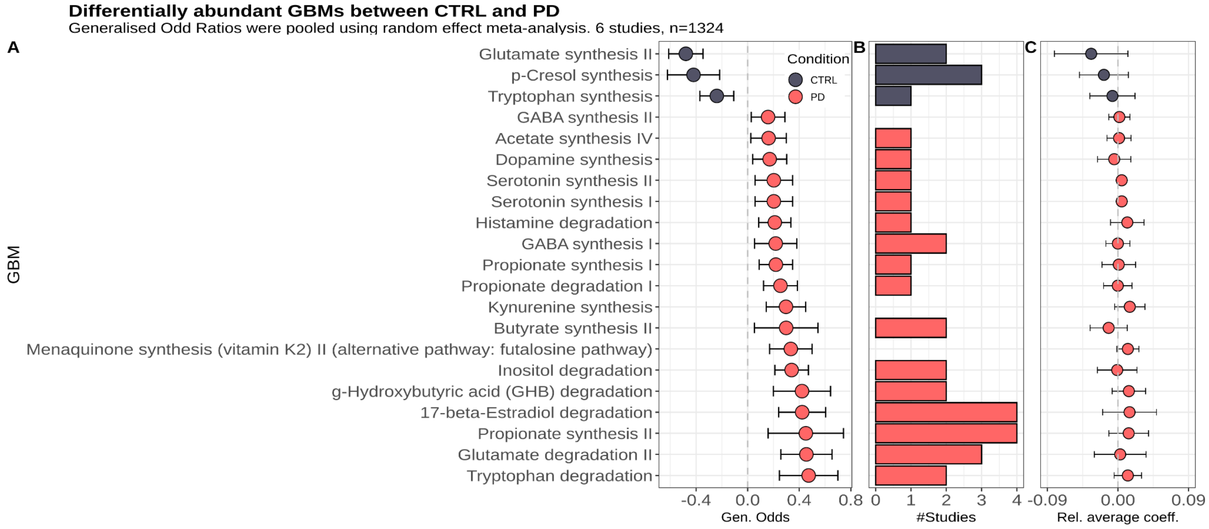


***Fig S19 | Proportion of bacterial pathways potentially confounded by covariates.*** *Heatmap showing the proportion of features associated with PD potentially confounded by sex and age (A), general medications (B), and PD medications (C). KEGG pathways, KEGG modules, and KO are related to the functionalities reported in Fig 6. Grey tiles indicate the absence of specific KEGG modules, within a given pathway, associated with PD. All microbiome features, including both taxa and functions, potentially confounded are reported in Supplementary Data 10-13.*

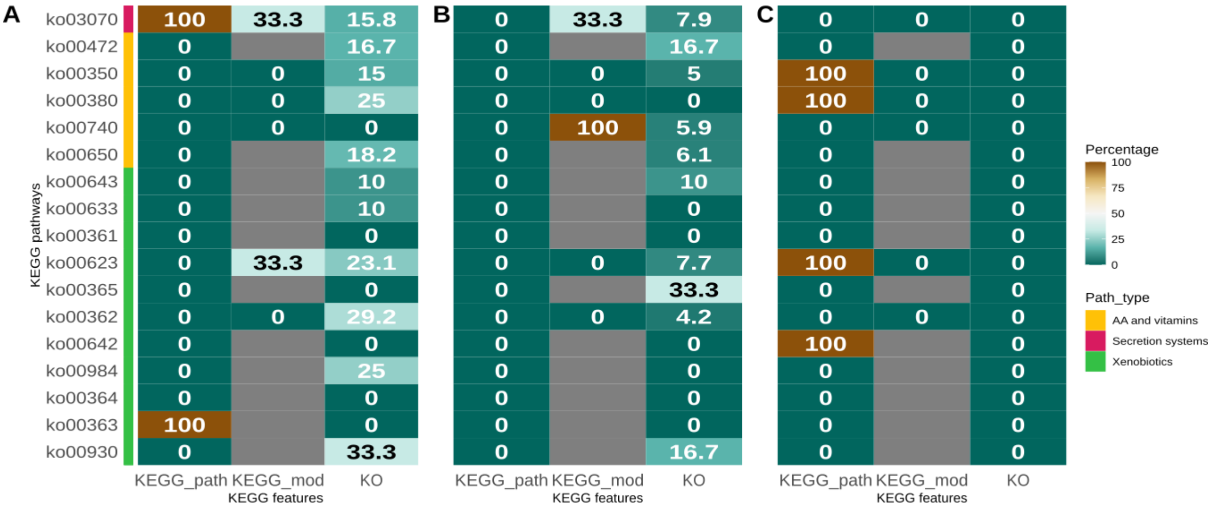
